# Neutrophils exposed to a cholesterol metabolite secrete extracellular vesicles that promote epithelial-mesenchymal transition and stemness in breast cancer cells

**DOI:** 10.1101/2024.08.02.606061

**Authors:** Natalia Krawczynska, Yu Wang, Ki Lim, Anasuya Das Gupta, Adam Lenczowski, Marwan Abughazaleh, Shruti V. Bendre, Lara I. Kockaya, Claire P. Schane, Yifan Fei, Alvaro G Hernandez, Jenny Drnevich, Jefferson Chan, Lawrence W. Dobrucki, Marni D. Boppart, Julie Ostrander, Erik R. Nelson

## Abstract

Small extracellular vesicles (sEVs) are emerging as critical mediators of intercellular communication in the tumor microenvironment (TME). Here, we investigate the mechanisms by which sEVs derived from neutrophils treated with the cholesterol metabolite, 27-hydroxycholesterol (27HC), influence breast cancer progression. sEVs released from 27HC treated neutrophils enhance epithelial-mesenchymal transition (EMT) and stem-like properties in breast cancer cells, resulting in loss of adherence, increased migratory capacity and resistance to cytotoxic chemotherapy. Decreased microRNAs (miRs) within the sEVs resulted in activation of the WNT/β-catenin signaling pathway in recipient cells and suggest that this may be a predominant pathway for stem-like phenotype and EMT. Our findings underscore a novel mechanism by which 27HC-modulated neutrophils contribute to breast cancer pathophysiology through EV-mediated intercellular communication, suggesting potential therapeutic targets in cancer treatment.

## INTRODUCTION

Despite significant advances in diagnosis and treatment, metastatic disease accounts for more than 90% of breast cancer-related deaths. Breast cancer is the most diagnosed cancer and the leading cause of cancer-related death in women world-wide^1,2^. Triple-negative breast cancer (TNBC) accounts for approximately 15–20% of breast cancer cases^3^ and 10–20% of invasive breast cancers^4^. TNBC is defined as a type of breast cancer with negative expression of estrogen (ER), progesterone (PR), and human epidermal growth factor receptor-2 (HER2), and thus is not sensitive to endocrine therapy. Therefore, with the exception of BRCA pathogenic variant carriers or the use of immune checkpoint blockers in the adjuvant setting, surgery followed by combined chemotherapy and radiation interventions are the main treatments for this type of breast cancer^5–9^. For those reasons, identifying new therapeutic targets for the successful treatment of metastatic breast cancer, especially for the TNBC subtype, is urgent.

Cholesterol is essential for maintaining the normal function of cells; it influences cell membrane integrity and lipid metabolism^10^. Cholesterol can be supplemented from the diet or synthesized through the *de novo* biosynthesis pathway. It is catabolized through two major bile acid synthesis pathways initiated by rate limiting enzymes CYP7A1 and CYP27A1, respectively^11^. As it is metabolized into bile acids, several oxysterol intermediates are formed^12^. 27-hydroxycholesterol (27HC) is the most abundant oxysterol in human circulation^13,14^ and works as a selective estrogen receptor modulator (SERM) and liver X receptor modulator (SLXRM)^15–17^. Both cholesterol and 27HC play important roles in regulating cellular function^17–19^. Cholesterol upregulation has been established as a risk factor for breast cancer and recurrence^12,20–22^. Whether circulating concentrations of 27HC is a risk or prognostic factor is less clear, but likely depends on breast tumor subtype, menopausal status, racial and ethnic group^13,23,24^. Since CYP27A1, the enzyme that synthesizes 27HC, is highly expressed in myeloid immune cells, circulating 27HC may not be reflective of tumor biology (risk/prognosis), but rather local concentrations being predictive. Indeed, in preclinical models, the genetic deletion of CYP27A1 only in myeloid immune cells was sufficient to impair tumor growth and metastasis of mammary and ovarian tumors^25,26^. Breast tumors have higher 27HC concentrations compared to adjacent normal and healthy breast tissue^18^. In addition, higher mRNA expression of the enzyme responsible for the catabolism of 27HC, CYP7B1 has been associated with better overall patient survival rates in breast cancer patients^27^.

27HC has been found to robustly increase metastasis in murine models of mammary cancer in a manner requiring myeloid immune cells^26,28^. At each step of cancer development, from initiation through promotion and metastatic progression, cancer cells have to escape from immune surveillance to survive in the host system^29^. Mechanistically, 27HC works through the LXR to shift myeloid immune cells into an immune-suppressive phenotype, impairing T cell expansion and function^25,26,30^. For mammary metastasis to the lung in mice, the neutrophil myeloid cell type (CD11B^+^;Ly6G^+^ cells) was found to mediate the pro-metastatic effects of 27HC^28^. The most abundant leukocytes in the blood, neutrophils have been underappreciated in cancer development. However, in the last decade, several seminal studies have implicated neutrophils in breast cancer progression and metastasis^31–39^. Neutrophils can infiltrate both the tumor microenvironment (TME) and pre-metastatic niche^40,41^. Due to their plasticity and heterogeneity, neutrophils have been reported to be anti-tumor and pro-tumor immune cells^40,42,43^. Interestingly, treatment with exogenous 27HC was found to increase the number of neutrophils within tumors and metastatic lesions of preclinical murine models of mammary cancer^19^. Subsequent work found that 27HC shifts myeloid cells, such as neutrophils, into a highly immune-suppressive phenotype by modulating the activity of the LXR^26,30^.

While investigating the mechanisms by which 27HC regulates neutrophils and metastasis, we found that 27HC increased the secretion of small extracellular vesicles (sEVs, also known as exosomes) from neutrophils^44^. sEVs from neutrophils treated with 27HC stimulated tumor growth and metastasis in murine models of TNBC^44^. However, it is not known how these 27HC-derived sEVs stimulate metastasis.

sEVs are lipid-bilayer membrane-enclosed vesicles that are heterogeneous in size, 30nm-150nm, and released from all cell types. sEVs serve as a method of intercellular communication between different cell types, transferring a complex mixture of cargo including DNA, RNA, proteins, metabolites, and even mitochondria under some conditions ^45,46^. There are a limited number of studies focusing on sEVs from neutrophils in cancer progression^47^. For example, sEVs from neutrophils have been implicated in reprogramming the TME and can promote both tumor progression and regression^48^. They have also been implicated in the metastatic process, mediated via nicotine activation, in lung^49^ and breast^50^ cancers. Thus, how the cargo of sEVs is altered by different conditions and how neutrophil-derived sEVs influence tumor pathophysiology are critical to our understanding of tumor progression, and ultimately developing therapies around them.

Here, we describe our findings that sEVs from 27HC-treated neutrophils led to mammary cancer cells losing their adherent properties. MicroRNAs (miRs) in normal sEVs from neutrophils inhibited the WNT/β-catenin pathway in recipient cancer cells, a regulatory mechanism which was lost in sEVs from 27HC-treated neutrophils. Consequently, cells receiving sEVs from 27HC-treated neutrophils gained a stem-like phenotype and underwent epithelial-mesenchymal transition (EMT), the consequences of which may lead to increased metastasis and drug resistance.

## RESULTS

From henceforth, sEVs obtained from neutrophils (Ly6G positive cells isolated from bone marrow) treated with vehicle (DMSO) will be referred to as DMSO-sEVs, and, sEVs obtained from neutrophils treated with 27HC will be referred to as 27HC-sEVs.

### Chronic treatment of cancer cells with small extracellular vesicles from 27-hydroxycholesterol treated neutrophils (27HC-sEVs) leads to loss of adherence

We have previously shown that treatment using 27HC-sEVs promoted breast cancer tumor growth and metastasis in animal models^44^. It was also found that cancer cells were able to take up 27HC-sEVs, suggesting that they may be a primary target^44^. To start elucidating the mechanisms by which 27HC-sEVs promote tumor progression, we performed an *in vitro* experiment, where we constantly exposed cancer cells to sEVs from neutrophils, mimicking the chronic exposure one would expect in the TME. We conducted this experiment in two independent TNBC murine cell lines: 4T1 and EMT6. The 4T1 cell line was derived from 410.4 tumor originating from the spontaneous mammary tumor of a BALB/c mouse, which closely resembles stage IV of human breast cancer^51^. The EMT6 cell line is a mammary murine carcinoma derived from BALB/c mouse^52^. Both lines are metastatic, with lesions found primarily in the lungs.

Cancer cells were treated with sEVs isolated from conditioned media of neutrophils treated with either vehicle (DMSO) or 27HC. To our surprise, the cancer cells exposed to 27HC-sEVs started to detach from the plates (**Fig. 1A-B**). The kinetics of detachment varied with 4T1 seeing peek detachment at 72h and EMT6 at 48h. To quantify this phenomenon, we determined the fraction of detached and attached cells using trypan blue exclusion to assess only live cells. There was a statistically significant difference between detached and attached 4T1 cells treated with 27HC-sEVs compared to DMSO-sEVs (**Fig. 1C**). Both live and dead cells were found attached and de-attached, for both treatment groups (**Fig. 1D**). When comparing adherent to non-adherent cells from 27HC-sEVs treatment, there was a slight increase in the proportion of dead cells in the non-adherent fraction, but this did not account for a large increase in non-adherent cells after this treatment (**Fig. 1C-D**). Moreover, when comparing live-adherent to live-non-adherent cells in our treatment groups, we found that treatment with 27HC-sEVs increased the live-non-adherent subpopulation (p=0.0617; **Fig. 1D-E**). Importantly, the loss of adherence was conserved in a second cell line examined (EMT6, **Fig. 1B, F-H**). For EMT6 cells, the non-adherent fractions for both treatments had increased dead cells compared to the adherent, providing further evidence that the detachment effect of 27HC-sEVs was not due to an overall increase in cellular death (**Fig. 1G**). The fraction of live-non-adherent cells was significantly larger in EMT6 cells treated with 27HC-sEVs (**Fig. 1G-H**).

**Figure 1.**
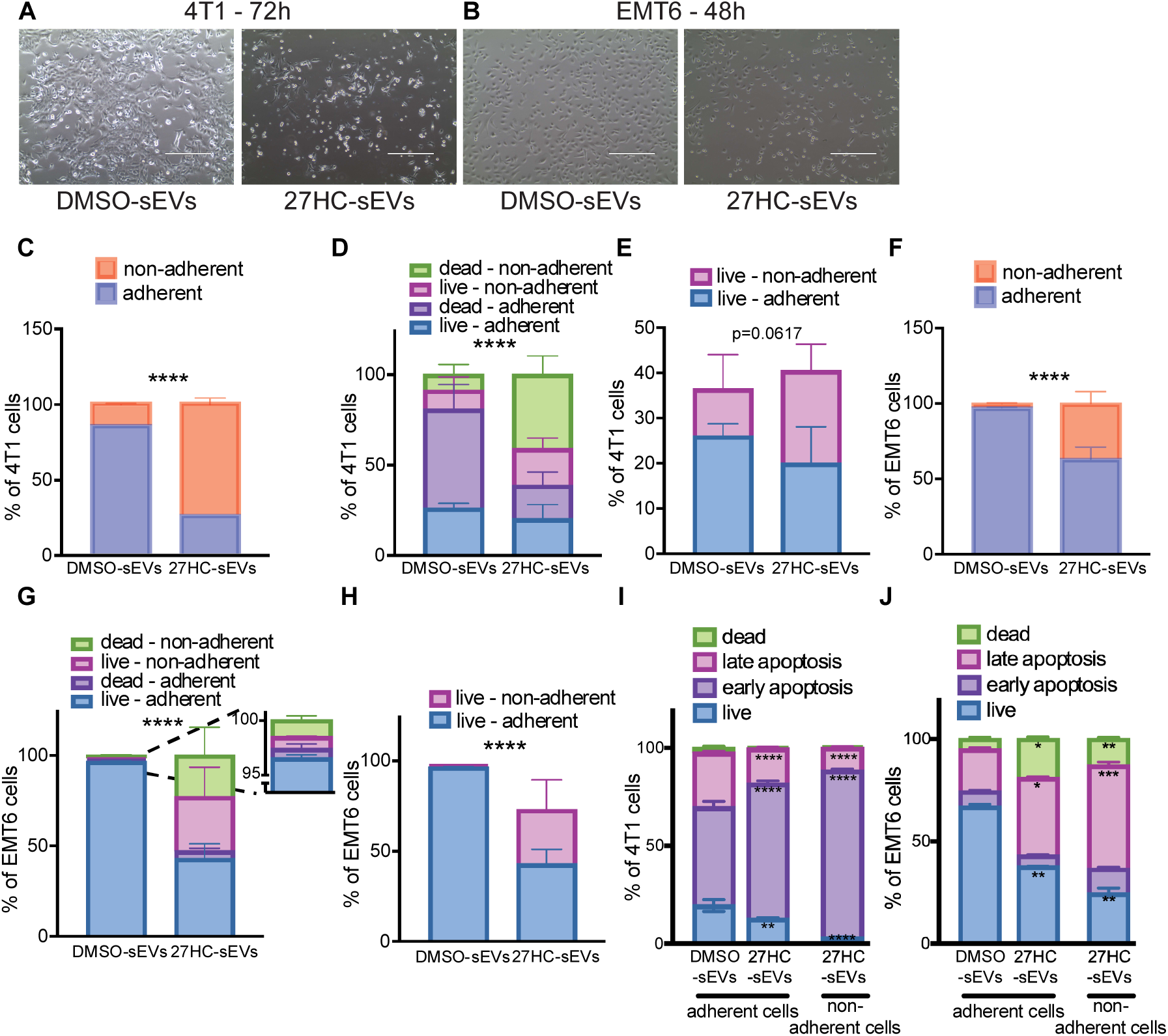
sEVs from 27-hydroxycholesterol treated neutrophils promote loss of adherence in breast cancer cells. Representative pictures of 4T1 **(A)** and EMT6 **(B)** cancer cells after exposure to DMSO-sEVs or 27HS-sEVs for 72h or 48h, respectively. n=3 **(C)** Quantification of attached vs. detached 4T1 cells 72h of the treatments. Trypan blue staining and an automatic cell counter (Countless 3, Invitrogen) were used. Statistical analysis was performed using the Fisher exact test. n=3; ****P-value<0.0001. **(D)** Quantification of live and dead cells in the attached and detached subpopulation of 4T1 cells after 72h of the treatments. Trypan blue staining and an automatic cell counter (Countless 3, Invitrogen) were used. Statistical analysis was performed using the Fisher exact test. n=3; ****P-value<0.0001. **(E)** Quantification of live-adherent and live-non-adherent cells in the attached and detached subpopulation of 4T1 cells after 72h of the treatments. Trypan blue staining and an automatic cell counter (Countless 3, Invitrogen) were used. Statistical analysis was performed using the Fisher exact test. n=3. **(F)** Quantification of attached vs. detached EMT6 cells 48h of the treatments. Trypan blue staining and an automatic cell counter (Countless 3, Invitrogen) were used. Statistical analysis was performed using the Fisher exact test. n=3; ****P-value<0.0001. **(G)** Quantification of live and dead cells in the attached and detached subpopulation of EMT6 cells after 48h of the treatments. Trypan blue staining and an automatic cell counter (Countless 3, Invitrogen) were used. Statistical analysis was performed using the Fisher exact test. n=3; ****P-value<0.0001. **(H)** Quantification of live-adherent and live-non-adherent cells in the attached and detached subpopulation of EMT6 cells after 48h of the treatments. Trypan blue staining and an automatic cell counter (Countless 3, Invitrogen) were used. Statistical analysis was performed using the Fisher exact test. n=3; ****P-value<0.0001. Quantification of flow cytometry analysis of 4T1 **(I)** and EMT6 **(J)** cells after 72h and 48h of the treatments, respectively, stained with annexin V and PI for live-dead cells evaluation. Cells were divided into 4 subpopulations: live cell (Annexin V-/PI-), early apoptotic cell (Annexin V+/PI-), late apoptotic cell (Annexin V+/PI+), and dead cell (Annexin V-/PI+). Statistical analyses were performed using one-way ANOVA. n=3; ****P-value<0.0001, ***P-value<0.001, **P-value<0.01, *P-value<0.05. All data are presented as mean+/-SEM.

To further rule-out whether 27HC-sEVs were inducing cellular death resulting in detachment, we evaluated apoptosis by staining with Annexin V-FITC and propidium iodide (PI) (gating strategy is presented in **Supplementary Fig. 1A-B**). Annexin V binds to phosphatidylserine, a membrane lipid that becomes accessible on the outer layer during cell membrane disruption. PI binds to DNA after entering the cell through an already disrupted membrane. It is important to note that although there were some non-adherent cells found in the media after treatment with DMSO-sEVs, the total number was far lower and too low to process with Annexin V / PI staining. Treatment with 27HC-sEVs did increase cells in the early and late apoptosis, in both remaining adherent and non-adherent cells (**Fig. 1I**). A similar, however distinct effect was observed in the EMT6 cell line after 48h of the treatments, with a significant subpopulation of live cells (Annexin V-/PI-) after treatment with 27HC-sEVs in comparison to DMSO-sEVs (**Fig. 1J**). For both cell types, there was also a significant fraction of live, non-adherent cells (**Fig. 1I-J**). Collectively, these data indicate that 27HC-sEVs impact cancer cells by promoting loss of adherence, resulting in the majority entering apoptosis but with a significant population of surviving cells. This mirrors an accepted paradigm *in vivo* where the majority of cells intravasating do not survive, but a small population does which goes on to form metastatic lesions^53^.

### Elevated 27HC modulates the miR content of small extracellular vesicles isolated from neutrophils

Using simultaneous label-free autofluorescence multiharmonic (SLAM) microscopy, we previously found that sEVs from 27HC-treated neutrophils have a decreased FAD:(FAD+NAD(P)H) ratio compared with sEVs from DMSO-treated neutrophils^44^. This strongly suggested that there is a difference in the cargo of 27HC-sEVs compared to DMSO-sEVs. Therefore, in order to investigate the potential mechanisms behind loss of adherence after exposure to 27HC-sEVs (**Fig. 1**) and ultimately metastatic progression^44^, we probed differences in the sEV molecular cargo. RNA is one of the major constituents of sEV cargo^54^. Therefore, we focused our efforts on RNA content.

Total RNA was isolated from DMSO-sEVs and 27HC-sEVs, using methods to also capture small amounts of RNA. First, we used Bioanalyzer 2100 (Agilent) and the Total RNA Pico kit detection method for fragment size, concentration, and quality verification. The RNA fragments were in the size range below 200 nucleotides (nt) (**Supplementary Fig. 2A-B**). The RNA integrity number (RIN) is based on the 28S:18S ratio, where a 2:1 ratio is considered the best RNA quality^55^. The observed relatively small or absent fragments of 18S and 28S in our tested samples highly suggest that the isolated RNA does not reflect the size distribution expected in a cell (as in, sEV RNA did not contain large RNA molecules, e.g. 18S and 28S). Thus, to evaluate the RNA species smaller than 200 nt, we performed a secondary analysis using the DNF-470-22 - Small RNA kit on an AATI Fragment Analyzer (Agilent). For all tested samples of total RNA isolated from sEVs, the detected fragments were in the range of 10-60nt, with the average peak at 18nt (**Supplementary Fig. 2C-D**). The calculated data revealed that, on average, 91.7% of isolated RNA qualified as microRNA (miR).

Therefore, we next focused on characterizing the miR cargo by performing small RNA-Seq on isolated sEVs from treated neutrophils, we powered our study with N=7 per group. In total, 1031 different miRs were detected in both treatment groups (**Supplementary Fig. 2E**). However, only 34 miRs were statistically differently expressed between the groups (**Fig. 2A**). Seven miRs were upregulated while 27 miRs were downregulated in 27HC-sEVs compared to DMSO-sEVs. We next validated our results in an independent experiment, employing the gold standard RT-qPCR method^56^. Since we could not assume that 27HC did not alter small RNAs typically used as internal controls, we decided to use an exogenous reference control by spiking – miR USP6 (Qiagen). As a further control, we assessed miR-103a-33p which was not found to be significantly regulated in the RNA-Seq, which was confirmed by qPCR (**Fig. 2B**, 1^st^ panel). Our results confirmed the changes in several of the differentially expressed miRs identified by small RNA-Seq of sEVs (**Fig. 2B**). To better understand the potential impact of the identified differentially expressed miRs in recipient cancer cells, we used three different bioinformatics pipelines (described in the Methods section with an overview summary in **Supplementary Fig. 3**) to predict targeted pathways. Even though we were able to detect statistical differences in miR expression between our treatment groups (**Fig. 2A**), when considered individually the prediction of potential downstream pathways for each miR was not informative. Therefore, future analysis was conducted using clusters of miRs.

**Figure 2.**
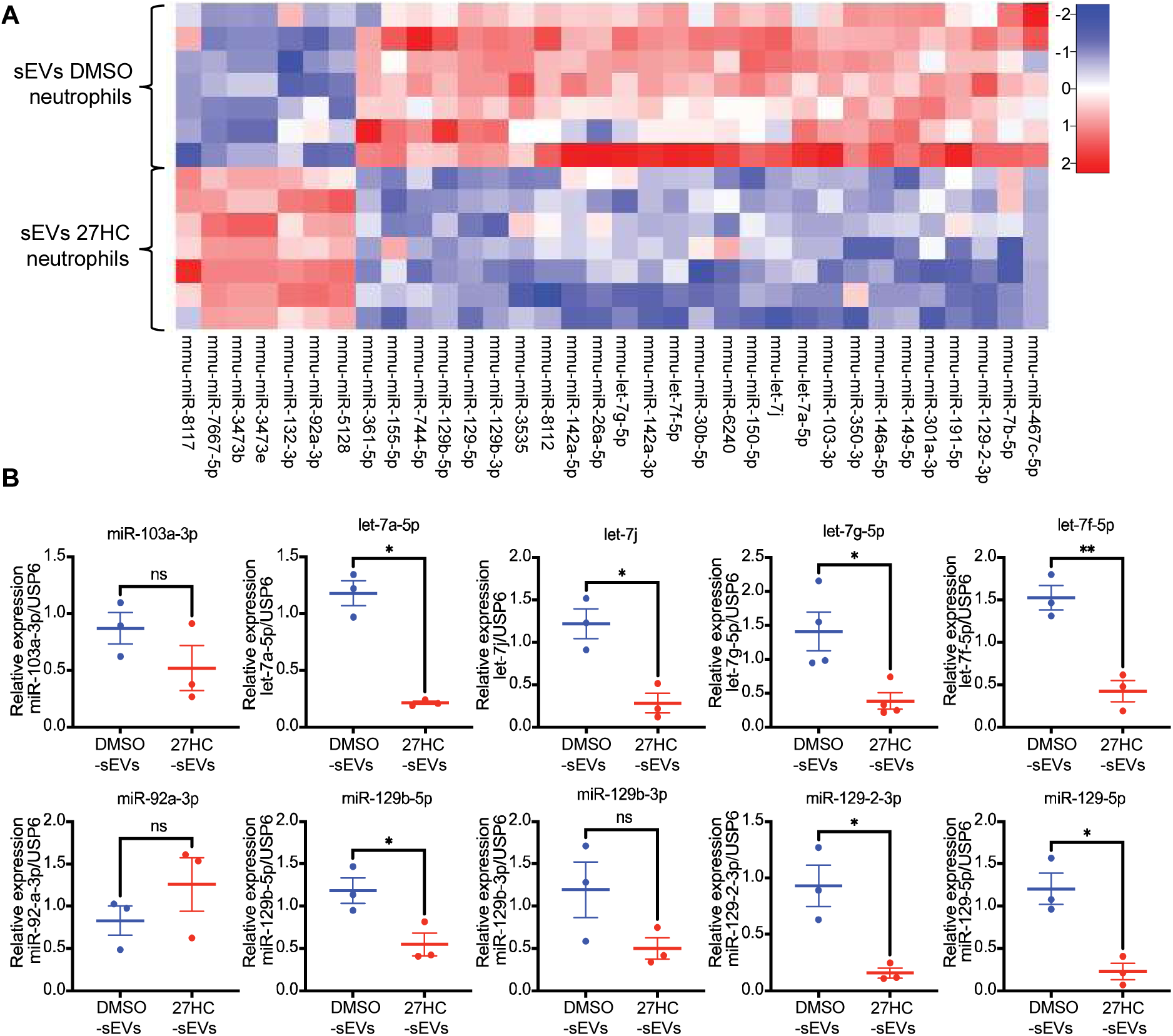
miR cargo of sEVs is altered after 27-hydroxycholesterol treatment of parental neutrophils. Murine neutrophils were treated with 27HC. Secreted sEVs were isolated from the media and processed for small RNA-Seq. **(A)** Heatmap of statistically significant, differently expressed microRNAs between sEVs isolated after treatment with either DMSO or 27HC. The False Discovery Method was used for statistical analysis. n=7/group **(B)** microRNA transcripts quantified by RT-qPCR from DMSO-sEVs and 27HC-sEVs. Statistical analyses were performed using Welch’s t-test. n=3; **P-value<0.01, *P-value<0.05. Data are presented as mean+/-SEM.

Based on this analysis, the downregulated miRs in 27HC-sEVs were predicted to have an impact on insulin signaling, response to hypoxia, GnRH signaling, PI3K-Akt-mTOR signaling, signaling pathways regulating pluripotency of stem cells, TGF-beta signaling, angiogenesis, EGF receptor signaling, WNT signaling, MAPK signaling pathways, response to amyloid-beta, oxidative stress response, and synaptic vesicle. Overall, 13 potential pathways that might be targeted by differentially expressed miRs were consistently implicated across the different bioinformatic approaches used (**Supplementary Table 1**). Gaining stem-like properties (e.g. pluripotency) by cancer cells was detected in two bioinformatic pipelines and caught our attention given our observations that 27HC-sEVs stimulated loss of adherence. Cancer stem cells (CSCs) are a small subset of cancer cells that are characterized by a low proliferation index with asymmetric division and self-renewal capabilities and the ability to survive in low attachment conditions^57–59^, they are considered the main cause of metastatic recurrence and drug resistance^60^. There is no universal marker to distinguish CSCs from other cancerous and non-cancerous cells. Breast cancer stem cells (BCSC) are generally characterized as CD44^+^/CD24^-/low^, and ALDH1A1^+^ ^60–62^. Gaining pluripotency or stem-like phenotypes may be driven by multiple pathways^57^. Our analysis of differential miRs predicted several downstream pathways that are implicated in driving pluripotency and ‘stemness’. Specifically, the PI3K-Akt-mTOR, TGF-beta, WNT, MAPK, and hypoxia signaling pathways have all been previously implicated in breast cancer progression, metastasis, and drug resistance^57,58,63^.

### 27-hydroxycholesterol also changes miR content of neutrophils, the cellular source of sEVs

It has been reported that the cargo of sEVs can be selectively loaded, as in their contents do not necessarily reflect those of their parental cells^64–66^. To better understand the origins of the differentially expressed miRs in sEVs, we simultaneously quantified select miRs in the parental neutrophils (**Supplementary Fig. 4A, 4C**) and resulting sEVs (**Supplementary Fig. 4B, 4D**). After 24h of treatment, miR expression levels in neutrophils had shifted but were not yet statistically significant (**Supplementary Fig. 4A**). At this timepoint, no changes were observed in sEVs apart from let-7j which was slightly upregulated (**Supplementary Fig. 4A**). Since many of the miRs of interest were downregulated by 27HC and miR turnover may take more than 24h, we next assessed a 48h timepoint. At this timepoint, the majority of miRs probed were significantly regulated by 27HC in neutrophils (**Supplementary Fig. 4C**). The differences observed in neutrophils 48h post-treatment were also reflected in the cargo of resulting sEVs (**Supplementary Fig. 4D**).

These results, comparing miR expression in neutrophils to sEVs, strongly suggest that 27HC modulates miR content within neutrophils themselves and that the miR cargo in sEVs is reflective of this change. However, the process may consist of multiple steps, and although our kinetic data would suggest the regulation of miRs primarily at the cellular level, we cannot rule out selective loading into sEVs. The mechanisms by which 27HC modulates miRs in the neutrophil or how they are loaded into sEVs require further investigation.

### sEVs from 27HC treated neutrophils disrupt the WNT/β-catenin pathway and impact stem-like properties in recipient cancer cells

Our bioinformatic analyses predicted 13 biologic pathways to be regulated in recipient cells by the miRs differentially expressed in 27HC-sEVs. To better understand the mechanisms of how 27HC-sEVs promoted metastatic progression, we performed a nonbiased, bulk RNA-Seq on 4T1 mammary cancer cell after exposure to DMSO-sEVs or 27HC-sEVs. To best capture dynamic regulation by sEVs miR cargo, we evaluated two-time points and powered these studies with five replicates per group. For cells exposed to 27HC-sEVs, we performed RNA-Seq on both cells that lost their adherence (non-adherent) and those that remained attached to the plate (adherent).

Using two-way ANOVA with an FDR cutoff of 0.05, 11,637 genes were found to be differentially expressed (DEGs) between the six groups (**Fig. 3A**). As illustrated by a Multidimensional Scaling Plot (MDS), exposure of 4T1 cells to 27HC-sEVs resulted in a unique transcriptional landscape compared to DMSO-sEVs; the most variance between groups being described by dimensions 1 and 2 (**Fig. 3B**). Within each treatment type, a shift in dimension 1 and 2 were observed through time (**Fig. 3B**). For those cells treated with 27HC-sEVs, both adherent and non-adherent cells clustered separately from those treated with DMSO-sEVs, particularly along dimension 1 (**Fig. 3B**). At the second timepoint (72h), the adherent cells treated with 27HC-sEVs started to more closely resemble nonadherent ones at the first timepoint (48h). Raw gene counts (GeneLevel_counts), and logarithmic counts per million reads of the trimmed mean of M-values (TMMnormalized_logCPM) are listed in **Supplementary Datasheet 1.**

**Figure 3.**
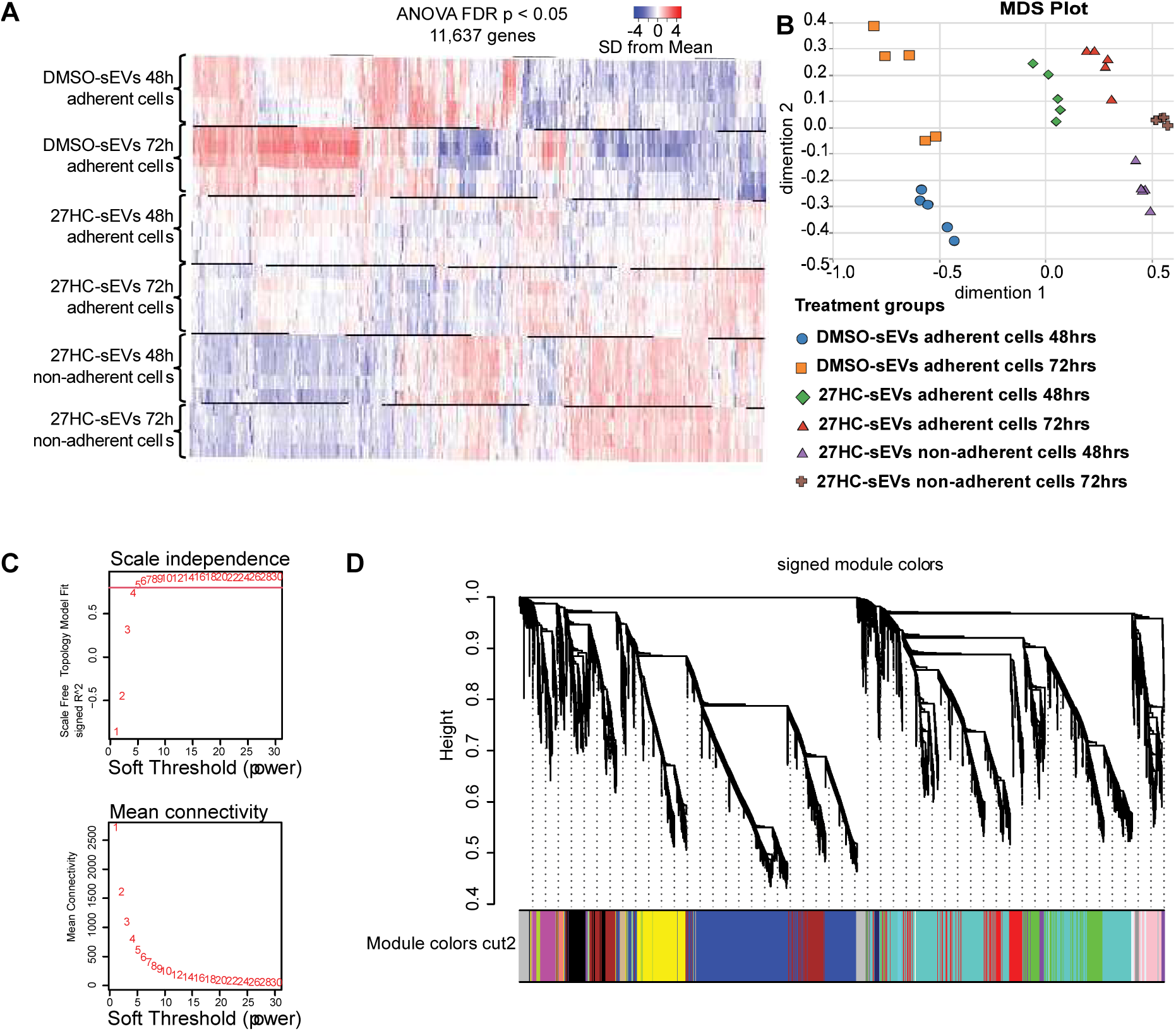
Transcriptome of cancer cells significantly changes upon treatment with 27HC-sEVs. Murine neutrophils were treated with DMSO or 27HC followed by sEV isolation from the conditioned media. 4T1 cancer cells were exposed to the sEVs for 48h or 72h. Both adherent and non-adherent cells were harvested and sent for bulk RNA-Seq. **(A)** Heatmap of DEGs between all tested groups. Statistical analyses were performed using two-way ANOVA, FDR <0.05. n=5 **(B)** MDS plot segregating and grouping samples based on dimensions 1 and 2. **(C)** WGCNA settings: The soft threshold power *β* settings are based on scale independence and mean connectivity. **(D)** WGCNA settings: a clustering diagram of the gene modules dendrogram by distinct colors. Each colored line represents a color-coded module containing highly connected genes.

The time, treatment, and, adherence variables in our experimental setup made it challenging to perform broad pathway analyses on the DEGs. Therefore, to focus our evaluation of 4T1 expression patterns following treatment with neutrophil-derived sEVs, we utilized weighted gene co-expression network analysis (WGCNA). The soft thresholding power *β* was set to five to ensure a correlation coefficient close to 0.8. Therefore, β = 5 (**Fig. 3C**) was used to generate a hierarchical clustering tree (**Fig. 3D**). A total of 16 different color-coded co-expression modules were identified. Each module is depicted in **Supplementary Fig. 5** and **Supplementary Datasheet 2**. Statistical analysis between treatment groups in specific modules (**Supplementary Table 2**) revealed the modules with the lowest P-value as Module 1 (co-downregulated genes, adj.P.Val = 6.80546234426343e-19) and Module 2 (co-upregulated genes, adj.P.Val = 1.55549927195221e-15) (**Fig. 4A-B, Supplementary Datasheet 3**). Corresponding heatmaps for each module are presented in **Fig. 4C-D**.

**Figure 4.**
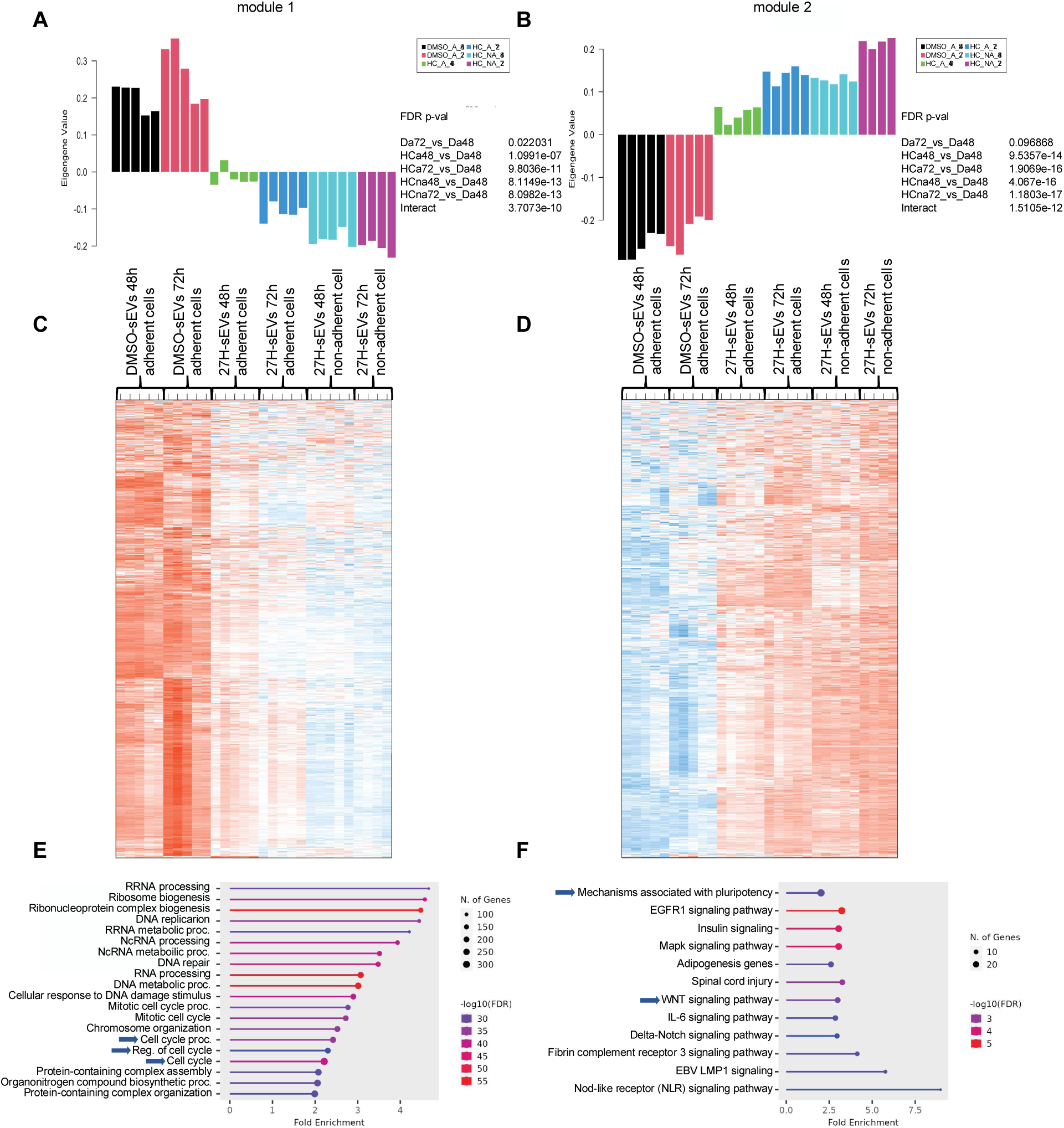
WGCNA analysis reveals several distinct modules of DEGs, the two most robust being comprised of genes associated with cell cycle, pluripotency and WNT signaling. RNA-Seq data from Fig. 3 was analyzed by weighted gene co-expression network analysis. The two most prominent modules depicted here, the remaining in Supplementary Fig. 5. (A) Module 1 co-downregulated genes – samples segregation. (B) Module 2 co-upregulated genes – samples segregation. (C) Module 1 – heatmap of co-downregulated genes between the treatment groups. (D) Module 2 – heatmap of co-upregulated genes between the treatment groups. (E) Module 1 and Module 2 (F) gene set enrichment analysis based on Shiny 0.8 software.

Gene-set enrichment analysis of Module 1 and Module 2 hub genes was performed using Shiny 0.8^67^. Enrichment of genes associated with cell cycle was found in Module 1, with 27HC-sEVs resulting in their down-regulation (**Fig. 4E**). Interestingly, in 4T1 cells treated 27HC-sEVs, there was significant enrichment in genes associated with the WNT signaling and the pluripotency (Module 2, upregulated by 27HC-sEVs, **Fig. 4F**). Collectively, these analyses suggest that 27HC-sEVs downregulate the cell cycle and division process while increasing WNT signaling and pluripotency. It is recognized that decreased cell-cycle is a characteristic feature of cancer stem cells^68^ and that WNT/β-catenin signaling is an important driver of pluripotency^57^. These data provide a potential mechanism for our observations that exposure of cancer cells to 27HC-sEVs lose their adherent properties, another common feature of pluripotent or stem-like cells^69^.

### sEVs from 27HC-treated neutrophils alter the expression of several genes in the WNT/β-catenin pathway

We first focused on the WNT/β-catenin pathway, as it has been implicated in EMT, driving stem-like states and loss of adherence^69–71^. Moreover, this pathway was also predicted to be regulated by miRs identified in our small RNA-Seq of sEVs (**Fig. 2**, **Supplementary Table 1).** Therefore, we performed independent experiments examining both 4T1 and EMT6 cells exposed to sEVs for two different timeframes. Gene expression analysis of several genes involved in the WNT/β-catenin pathway found that many were altered, with the largest differences observed 27HC-sEVs treated cells that had lost their adherence (**Supplementary Fig. 6**). The expression of: WNT5A, WNT9B, WNT2B, FRZ3, FRZ6, CTNNB1, APC, AXIN2, TCF7 was increased. At the same time, GSK3b was decreased in 4T1 cells after 72h (**Supplementary Fig. 6A**) and 96h (**Supplementary Fig. 6B**) compared to cells treated with DMSO-sEVs. A similar pattern was observed in the EMT6 cell line after 48h (**Supplementary Fig. 6C**) and 72h (**Supplementary Fig. 6D**).

Glycogen synthase kinase-3b (GSK3b) negatively regulates the WNT/β-catenin pathway by promoting the phosphorylation and subsequent proteasomal degradation of β-catenin^72^. When GSK3b is inhibited, β-catenin accumulates in the cytoplasm and is translocated to the nucleus, where it functions as a co-transcriptional activator for T-cell factor/lymphoid enhancer factor (TCF/LEF) transcription factors, upregulating transcription of c-myc, WNT genes, and SOX2^72,73^. In our case, GSK3b was downregulated by 27HC-sEVs, while β-catenin (CTNNβ1) was upregulated, reinforcing this signaling loop (**Supplementary Fig. 6**). In addition, other genes activating the pathway were upregulated by 27HC-sEVs. Collectively, these data indicate that 27HC-sEVs robustly increase WNT signaling in mammary cancer cells, suggesting that this may be a predominant pathway in the stemness and EMT phenotypes observed.

### sEVs from 27HC-treated neutrophils promote pluripotency and stemness in cancer cells

The impact of transferred miRs by 27HC-sEVs to recipient cells on the pluripotency of stem cells was concluded based on bioinformatic analysis of potential miR-targeted pathways (see Results section *Elevated 27HC modulates the miR content of small extracellular vesicles isolated from neutrophils*). In addition, analysis of RNA-Seq from 4T1 cells implicated pathways involved in pluripotency (**Fig. 4F**). Therefore, we assessed the expression of transcription factors known to play central roles in driving key transcriptional programs related to pluripotency and stemness: SOX2, OCT4, and NANOG^74^. As anticipated, the expression of SOX2, OCT4, and NANOG were upregulated in the resulting non-adherent 4T1 cells after treatment with 27HC-sEVs, 72h after exposure (**Supplementary Fig. 7A**). Since we observed loss of adherence in EMT6 cells sooner (48h timepoint; **Fig. 1**), we examined this timepoint for changes in these transcription factors for EMT6. Similar to our findings in 4T1 cells, 27HC-sEVs increased the expression of SOX2, NANOG, and OCT4 (**Supplementary Fig. 7B**). For both cell types, the remaining adherent cells post 27HC-sEVs treatment also had higher expression of these transcription factors compared to DMSO-sEVs, resembling an intermediate state (**Supplementary Fig. 7A-B**). Gaining pluripotent ability (by overexpression SOX2, OCT4, and NANOG) by cancer cells can give rise to multiple types of cells within the tumor, contributing to tumor heterogeneity and chemoresistance^60,68,74^.

Proliferative capacity is often low or reduced in stem-like cells^59^. As expected, we observed significantly reduced expression of the proliferation marker, ki67, through time in 27HC-sEVs treated 4T1 and EMT6 cells (**Supplementary Fig. 7C-D**). The reduction in ki67 was most pronounced in non-adherent populations, which emerged at 72h or 48h post-treatment for 4T1 and EMT6 cells, respectively. This suggests that 27HC-sEVs from neutrophils were indeed promoting stemness.

Cancer stem cells in human breast tumors have been characterized as Lin^neg^/CD24^neg^ ^to^ ^lo^/CD44^+^ phenotype^75^. Murine mammary cancer stem cells have also been described to highly express CD44 and lose expression of CD24^76^. CD44 is one of the regulators of self-renewal, tumor initiation and metastatic progression, and chemoresistance properties of cancer stem cells^77^. On the other hand, CD24 is a negative regulator of CXCR4; CD24^neg^ ^to^ ^lo^ phenotype upregulates CXCR4 and may give rise to increased migration activity, required for tumor initiation and metastatic progression^78^. A distinct breast cancer stem cell population was also identified and characterized by high activity of aldehyde dehydrogenase (ALDH), characterized by self-renewal capacity and multidrug resistance^61^. Moreover, the ALDH1A1 isoform is considered the main cancer stem cell-related isoform^79^. Based on the markers (ALDH, CD44, and CD24), breast cancer stem cells have been divided into subpopulations: epithelial-like, with ALDH1A1^+^ CD24^neg^ ^to^ ^lo^/CD44^+^, and mesenchymal-like, ALDH1A1^negative^ CD24^neg^ ^to^ ^lo^/CD44^+^^76,80^. Our flow cytometric analysis of 4T1 or EMT6 cells after exposure to 27HC-sEVs revealed a robust enrichment of the CD24^neg^/CD44^+^population (**Fig. 5A-B**). In addition, the expression of the ALDH1A1 marker decreased after 27HC-sEVs treatment in adherent and non-adherent 4T1 or EMT6 cells (**Fig. 5C-D**). The decrease in ALDH1A1 was most apparent in non-adherent cells at later timepoints (times after detachment was observed; **Supplementary Fig. 7E-F**). This indicates that after exposure to 27HC-sEVs, cancer cells are enriched in mesenchymal-like stem-like cancer cell subpopulations characterized by ALDH1A1^negative^ CD24^neg^ /CD44^+^.

**Figure 5.**
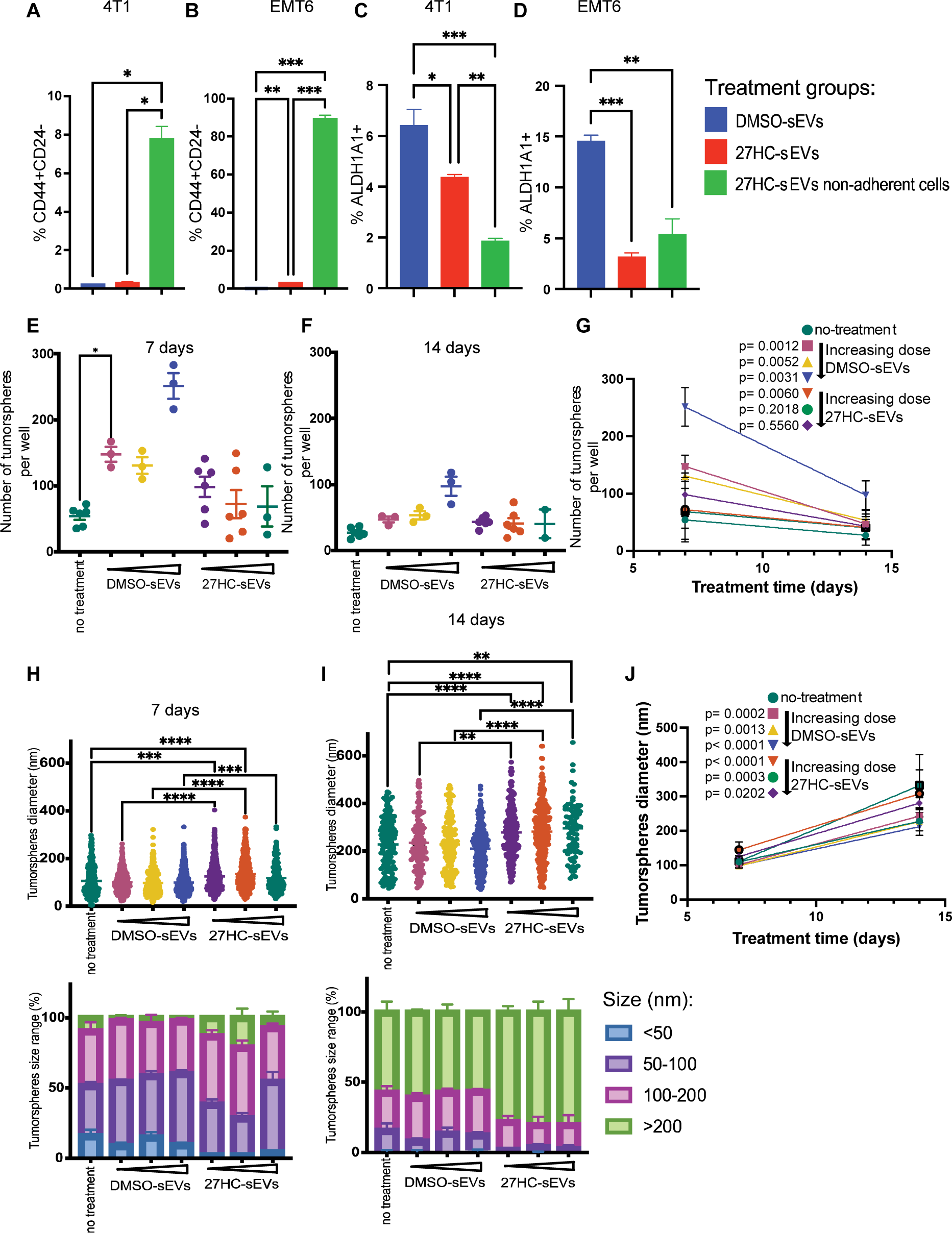
Treatment of mammary cancer cells with 27HC-sEVs from neutrophils results in a stem-like phenotype. Flow cytometry quantification of CD44^+^/CD24^-^ cells of **(A)** 4T1 after 72h and **(B)** EMT6 after 48h exposure to DMSO-sEVs or 27HC-sEVs. Statistical analyses were performed using one-way ANOVA followed by Tukey’s multiple comparison test n=3; ***P- value<0.001, **P-value<0.01, *P-value<0.05. Flow cytometry quantification of ALDH1A1^+^ cells of **(C)** 4T1 after 72h and **(D)** EMT6 after 48h of DMSO-sEVs and 27HC-sEVs treatments. Statistical analyses were performed using one-way ANOVA followed by Tukey’s multiple comparison test n=3; ****P-value<0.0001, ***P-value<0.001, **P-value<0.01, *P-value<0.05. Quantification of 4T1 tumorspheres after 7 days and **(E)** 14 days **(F)** of culture with different doses of DMSO-sEVs and 27HC-sEVs. Statistical analyses were performed using one-way ANOVA followed by Tukey’s multiple comparison test; *P-value<0.05. **(G)** Linear regression of 4T1 tumorsphere number after 7 days and 14 days of culture with different doses of DMSO-sEVs and 27HC-sEVs. Quantification of 4T1 tumorsphere diameters after 7 days **(H)** and 14 days **(I)** of culture with different doses of DMSO-sEVs and 27HC-sEVs. Statistical analyses were performed using one-way ANOVA followed by Tukey’s multiple comparison test; ****P-value<0.0001, ***P-value<0.001, **P- value<0.01. **(J)** Linear regression of 4T1 tumorsphere diameters from 7 days to 14 days of culture with different doses of DMSO-sEVs and 27HC-sEVs. All data are presented as mean+/-SEM.

A hallmark of stem-like cells is their ability to grow in an anchorage-independent fashion^81^. Therefore, we tested the ability of cancer cells to form tumorspheres in the presence of DMSO-sEVs or 27HC-sEVs. The assay was conducted for a total of 14 days with two assessment points – 7 and 14 days (**Fig. 5E-J, Supplementary Fig. 7G**). There was no statistical difference in the total number of tumorspheres between treatment groups (**Fig. 5E-F**) for both tested time points; with the exception of the no-treated group which was lower than the smallest dose of DMSO-sEVs neutrophils (7 days, **Fig. 5E**). However, there was a decrease in surviving tumorspheres between 7 days and 14 days of culture (**Fig. 5E-G**). The rate of this decrease was not as pronounced in tumorspheres treated with 27HC-sEVs (**Fig. 5G**). In contrast, the rate of tumorsphere loss was apparent in all doses of DMSO-sEVs examined (**Fig. 5G**).

Interestingly, we also observed larger diameters in 4T1 tumorspheres formed in the presence of 27HC-sEVs compared to DMSO-sEVs for both tested time points (**Fig. 5H-I**). Linear regression (**Fig. 5J**) revealed a dose-dependent growth of tumorspheres after 27HC-sEVs and DMSO-sEVs treatments.

Overall, sEVs from 27HC-treated neutrophils increase WNT signaling and upregulate the main transcription factors associated with pluripotency in mammary cancer cells. At the cellular level, mammary cancer cells treated with 27HC-sEVs gain a mesenchymal stem-like phenotype (ALDH1A1^negative^ CD24^neg^/CD44^+^) suggestive of epithelial-mesenchymal transition and, in consequence, lose their adherence.

### sEVs from 27HC treated neutrophils alter epithelial-mesenchymal transition in recipient cancer cells

Loss of adhesion is one of the hallmarks of metastasis^69^. In general, epithelial cancer cells lose dependence on integrin-mediated and cadherin-dependent interactions with extracellular matrix and cell-to-cell adhesion. For that reason, we started looking in detail into this phenomenon after sEVs treatments.

EMT is a reversible process that allows epithelial cells to adopt a mesenchymal state^80^. The “cadherin switch” is a characteristic movement assigned to the EMT. It is manifested by loss of E-cadherin (CDH-1) expression and increased N-cadherin (CDH-2) expression accompanied by different integrin changes. The downregulation of CDH-1 and upregulation of CDH-2 was observed in 4T1 cells after 72h and 96h treatment with 27HC-sEVs for both subpopulations: attached and non-adherent cells (**Fig. 6A-B**). The expression of genes associated with EMT were also altered in the EMT6 cell line (**Fig. 6C-D**). After 48h and 72h of treatment, both CDH-1 and CDH-2 were upregulated by 27HC-sEVs. This suggests that the EMT6 cells were in between epithelial and mesenchymal states, which is a common phenomenon for cancer cells^80^.

**Figure 6.**
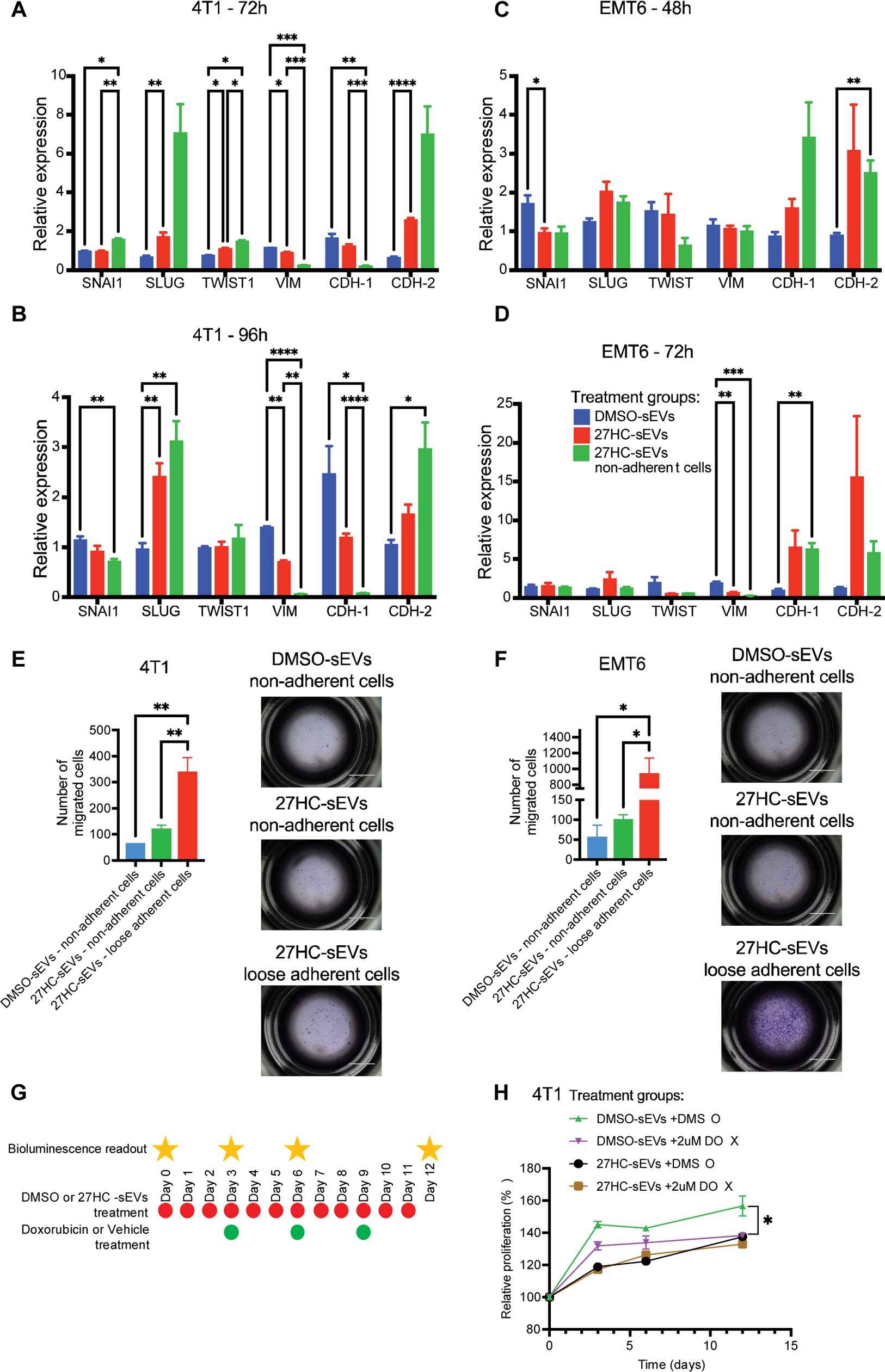
Treatment of mammary cancer cells with 27HC-sEVs from neutrophils results in an epithelial to mesenchymal transition (EMT) phenotype, increased migration and chemoresistance. EMT gene expression patterns in 4T1 cells after exposure to DMSO-sEVs or 27HC-sEVs for **(A)** 72h and **(B)** 96h. Statistical analyses were performed using one-way ANOVA followed by Tukey’s multiple comparison test, n=5-6; ****P-value<0.0001, ***P-value<0.001, **P-value<0.01, *P-value<0.05. EMT gene expression patterns in EMT6 cells after exposure to DMSO-sEVs or 27HC-sEVs for **(C)** 48h and **(D)** 72h. Statistical analyses were performed using one-way ANOVA followed by Tukey’s comparison test, n=5-6; ****P-value<0.0001, ***P-value<0.001, **P-value<0.01, *P-value<0.05. (**E**) Quantification of migrated 4T1 cells. 4T1 cells were cultured in the presence of either DMSO-sEVs or 27HC-sEVs for 72h prior to seeding onto a Boydon Chamber and allowing to migrate for 24h. Representative pictures for each treatment are presented to the right of the quantified data. (**F**) Quantification of migrated EMT6 cells. EMT6 cells were cultured in the presence of either DMSO-sEVs or 27HC-sEVs for 48h prior to seeding onto a Boydon Chamber and allowing to migrate for 24h. Representative pictures for each treatment are presented to the right of the quantified data. Statistical analyses were performed using one-way ANOVA followed by Tukey’s multiple comparison test, n=3; ****P-value<0.0001, ***P-value<0.001, **P-value<0.01, *P-value<0.05. (**G**) Schematic overview of 4T1 proliferation in 3D culture assay treatments and bioluminescence read. (**H**) Quantification of 4T1 3D proliferation assay over 12 days of treatments with DMSO-sEVs and 27HC-sEVs together with either doxorubicin or DMSO. Statistical analyses were performed using one-way ANOVA followed by Tukey’s multiple comparison test, n=3; *P-value<0.05. All data are presented as mean+/-SEM.

Stem and mesenchymal-like populations of cancer cells tend to be migratory^82,83^. To investigate the migratory ability of cancer cells after the sEVs treatments, we took detached subpopulations of cells after treatment with DMSO-sEVs or 27HC-sEVs and assessed their ability to migrate through a Boyden chamber. For both 4T1 and EMT6 cancer cells, we observed increased migration in cells treated with 27HC-sEVs, with the highest migration in loose-adherent (transitioning) 27HC-sEVs treated cells (**Fig. 6E-F**).

Another feature of stem cancer cells is resistance to cytotoxic chemotherapy. Drug resistance was assessed by 3D proliferation of 4T1 cells co-treated with DMSO-sEVs or 27HC-sEVs and doxorubicin (DOX) or vehicle (DMSO) through time (**Fig. 6G-H**). As expected, growth of 4T1 cells treated with DMSO-sEVs was inhibited by DOX treatment. In contrast, although they had a lower basal proliferation rate, 4T1 cells treated with 27HC-sEVs were not sensitive to DOX treatment. Thus, 27HC-sEVs impart resistance to cytotoxic chemotherapy, similar to what would be expected of CSC-like cells.

Collectively, these data strongly suggest that exposure to 27HC-sEVs initiates epithelial-mesenchymal transition and increased stemness. This allows cancer cells to detach from the plates, form persistent and larger tumorspheres, gain migratory ability and resist cytotoxic chemotherapies, all important features in the metastatic progression cascade.

### Let-7 miRs are a main regulatory component resulting in observed changes in recipient cancer cells

Even though we were able to detect statistical differences in the expression of several different miRs between our treatment groups (**Fig. 2A**), it was likely that no one miR was responsible for the phenotypes we observed. Therefore, future analysis was conducted when clusters of miRs were considered to impact gene expression.

We were able to identify two miR families that were downregulated in 27HC-sEVs: let-7 and miR-129s. Both miR families have been associated with aspects of WNT signaling and stemness. To test whether decreases in sEVs delivery of these miRs influenced cancer cells, we exposed our cancer cells to DMSO-sEVs or 27HC-sEVs, followed by miR quantification in the recipient cancer cells. miR-129 expression within 4T1 and EMT6 cells was below the detection limits of our qPCR approach. In both cell lines tested, let-7 miRs were decreased in 27HC-sEVs treated cells compared to DMSO-sEVs treated ones, with the largest decreases noted in the non-adherent cells (**Fig. 7**).

**Figure 7.**
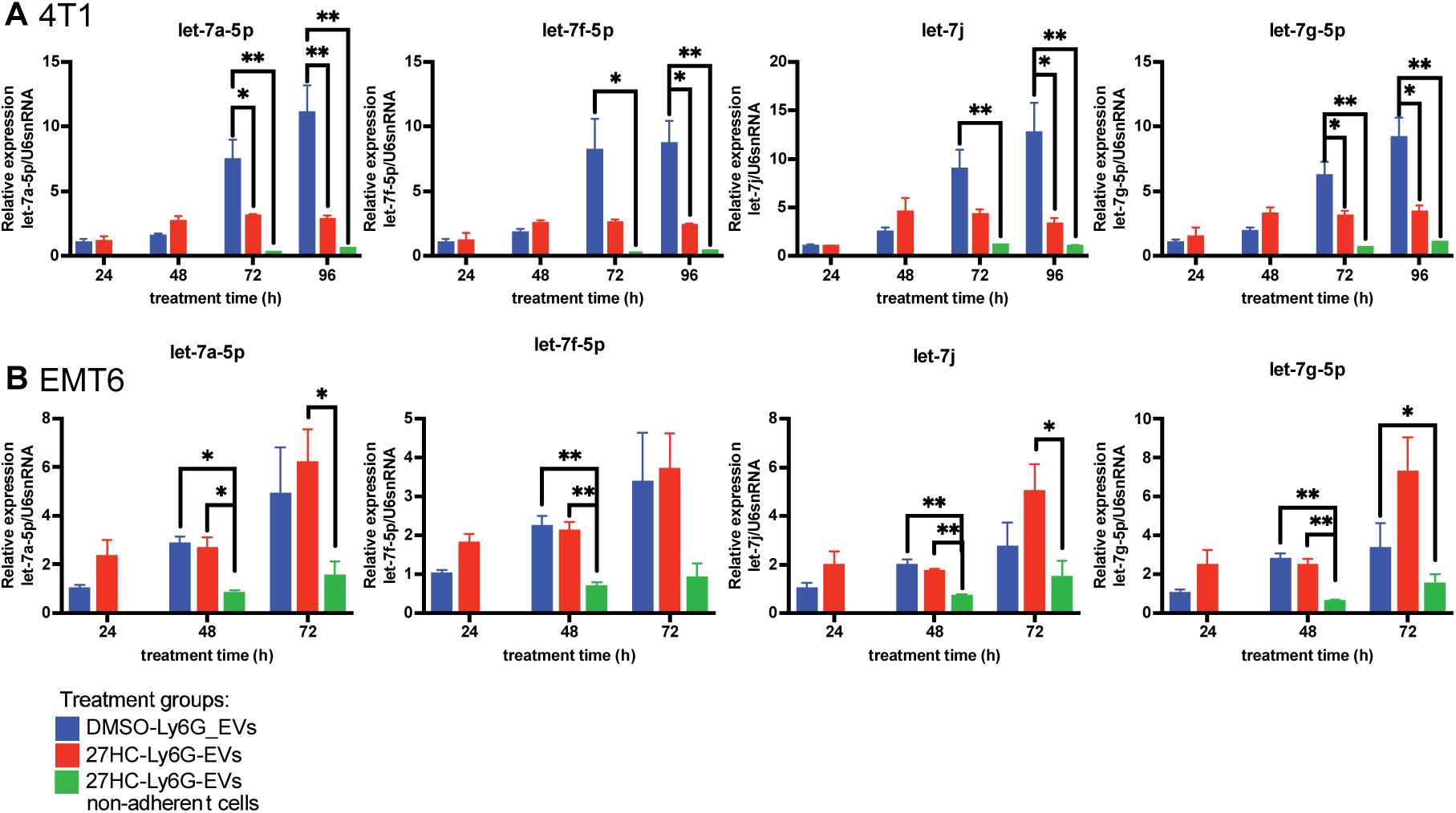
Expression of *Let-7 miRs decreases in cancer cells exposed to 27HC-sEVs.* Cancer cells were exposed for indicated timepoints with sEVs harvested from neutrophils treated with either DMSO or 27HC. **(A)** Expression of different let-7 miRs through time in 4T1 cells exposed to DMSO-sEVs or 27HC-sEVs. **(B)** Expression of different let-7 miRs through time in EMT6 cells exposed to DMSO-sEVs or 27HC-sEVs. Statistical analyses were performed using one-way ANOVA followed by Tukey’s multiple comparison test, n=3; ****P-value<0.0001, ***P-value<0.001, **P-value<0.01, *P-value<0.05. All data are presented as mean+/-SEM.

In addition, we engineered in-house sEVs derived from neutrophils with decreased levels of all the let-7’s that we identified as being downregulated after 27HC treatment. Neutrophils were treated with let-7a-5p, let-7f-5p, let-7g-5p, and let-7j inhibitors (antisense oligonucleotides; miRCURY LNA miRNA inhibitors), and sEVs were isolated. These sEVs were then used to treat cancer cells. First, we confirmed that let-7’s inhibitors work to reduce the expression of the expected let-7 miRs in neutrophils (**Fig.8A**) and sEVs (**Fig. 8B**). We compared the inhibitor group (let’7’s inh) to the negative control for the inhibitor treatment (NC), and untreated cells and sEVs (no treatment; NT).

**Figure 8.**
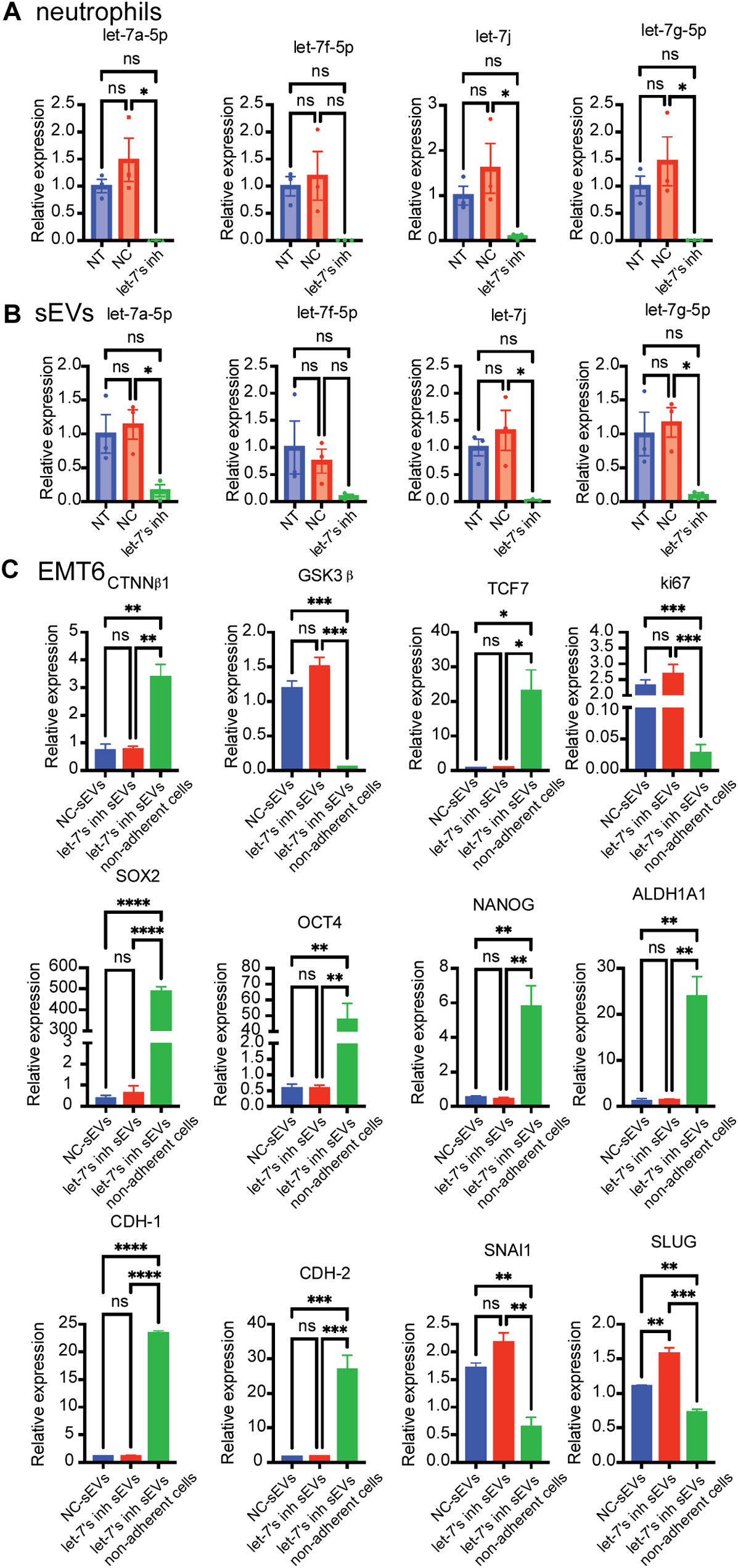
Decreased let-7 miRs in sEVs result in altered expression of genes associated with the WNT pathway and stemness in recipient cancer cells, mirroring the effects of 27HC-sEVs. Murine neutrophils were treated with a cocktail of antisense LNA oligos against different let-7 miRs (termed let-7 inh: let-7a-5p, let-7g-5p, let7f-5p, and let7j). **(A)** RT-qPCR quantification of let-7 miR in neutrophils after treatment with let-7 inh for 48h. NT - no treated cells, NC - negative control for miR inhibitors, let-7’s inh – the cocktail of 4 let-7’s inhibitors in the final concentration of 5nM. Statistical analyses were performed using one-way ANOVA followed by Tukey’s multiple comparison test, n=3; *P-value<0.05. **(B)** RT-qPCR quantification of let-7 miRs in sEVs secreted from neutrophils treated with let-7’s inhibitors for 48h. NT - no treated cells, NC - negative control for miR inhibitors, let-7’s inh – the cocktail of 4 let-7’s inhibitors in the final concentration of 5uM. Statistical analyses were performed using one-way ANOVA followed by Tukey’s multiple comparison test, n=3; *P-value<0.05. **(C)** Quantification of genes associated with the WNT pathway (CTNNB1, GSK3B, and TCF7), stem cells markers (ki67, SOX2, OCT4, NANOG, and ALDH1A1) or EMT (CDH-1, CDH-2, SNAI1, and SLUG) in EMT6 cells after 72h exposure to sEVs. NC -sEVs – sEVs were isolated from neutrophils treated with negative control for miR inhibitors; let-7’s inh_-sEVs. sEVs were isolated from neutrophils treated with the cocktail of 4 let-7 inhibitors at a final concentration of 5uM. Statistical analyses were performed using one-way ANOVA followed by Tukey’s multiple comparison test, n=3; ****P-value<0.0001, ***P-value<0.001, **P-value<0.01. All data are presented as mean+/-SEM.

Gene expression analysis revealed that let-7 inhibition phenocopied that of 27HC-sEVs in recipient cells (**Fig.8C**). As with 27HC-sEVs, WNT pathway genes (CTNNB1, GSKB, and TCF7) were upregulated. Moreover, cells gained a stem-like phenotype, based on downregulated ki67, and upregulated SOX2, OCT4, and NANOG. ALDH1A1 gene expression was upregulated, which was in contrast to our flow cytometry analysis of cells post-27HC-sEVs treatment (**Fig. 5D**). This suggests that after treatment with let-7 inhibitors sEVs EMT6 treated cells gain a more epithelial stem-like phenotype. Thus, it is likely that although the let-7 miR family is the predominant executor of the effects observed in 27HC-sEV treated cancer cells, other miRs or other cargo, in general, were fine-tuning responses.

Overall, the let-7 family is likely a major driver of cancer cell pluripotency and stem-like phenotype, after exposure to 27HCA-sEVs. This results in cancer cells that are more migratory and resistant to chemotherapy, signatures of cells thought to cause metastatic recurrence.

## DISCUSSION

EVs are crucial for maintaining regular cell-to-cell communication. They have been reported to play a significant role of messengers in tumor progression. It was previously reported that 27HC stimulates breast cancer metastasis in part by increasing the secretion of sEVs from neutrophils^28,44^. However, it was not known how these 27HC-derived sEVs stimulate metastasis. In this study, we have found that sEVs from 27HC-treated neutrophils impact cancer cells by promoting the loss of adherence (**Fig.1**) and initiating epithelial-mesenchymal transition (**Fig. 7**). Moreover, cancer cells gain pluripotency of stem cells and mesenchymal stem-like phenotype, characterized by ALDH1A1^negative^ CD24^neg^ ^to^ ^lo^/CD44^+^ (**Fig. 6**). Both EMT and stemness are driven by robust increases in WNT signaling, a pathway enriched by cancer cells exposed to 27HC-sEVs (**Fig. 3-4**). When probing the mechanism behind this, we found that sEVs from 27HC-treated neutrophils lost protective let-7 miRs (**Fig. 2**), resulting in decreased abundance of let-7 miRs in recipient cancer cells (**Fig. 8**). The decreased let-7 miR expression in sEVs was due to their regulation within the parental neutrophils, suggesting that it was not through selective cargo-loading (**Supplementary Fig. 4**). However, the mechanisms by which 27HC modulates miRs in the neutrophil or how they are loaded into sEVs are still unknown and need to be investigated.

miRs in sEVs from neutrophils inhibited the WNT pathway in recipient cancer cells, a regulatory mechanism lost in sEVs from 27HC-treated neutrophils. Consequently, cells receiving sEVs from 27HC-treated neutrophils gained a stem-like phenotype and underwent an epithelial-mesenchymal transition, the consequences of which may lead to increased metastasis and drug resistance. EMT endows cancer cells with increased motility and the ability to invade and disseminate to distant organs^69,80^. In addition, EMT is a reversible process, which can lead cancer cells to a hybrid epithelial/mesenchymal state. They can dynamically transition between those two stages, which allows them to gain properties from both phenotypes when needed^80^. Cancer stem cells can self-renew and differentiate, which contributes to tumor recurrence and heterogeneity^78,81^. Both EMT and stemness are strongly implicated in resistance to chemotherapy; in part due to their slower proliferation rate and higher expression of drug efflux pumps^83^. We found that 27HC-sEVs downregulated proliferation and promoted chemoresistance.

Understanding the dual processes of EMT and the acquisition of stemness in cancer cells is crucial for developing new therapeutic strategies to combat cancer progression, metastasis, and treatment resistance. The tumor microenvironment plays a significant role in regulating EMT and stemness. Key components driving those processes include cellular (immune cells, fibroblasts, adipocytes), signaling pathways (i.e. WNT ligands), hypoxic conditions, and extracellular matrix (ECM) remodeling^83,84^. However, little is known about how “the signal” is transduced and what exactly triggers those components that drive EMT and stemness. Here, we have found that when exposed to 27HC, neutrophils change their miR content and consequently release sEVs (“the signal”) with modulated molecular cargo. Those sEVs are taken by cancer cells and sensitize the WNT signaling pathway that is a known initiator of EMT and stemness^57–59,70–73^.

Targeting the 27HC pathway could offer a novel therapeutic approach in breast cancer patients^13^. Since 27HC influences the tumor and metastatic microenvironment by recruiting neutrophils and modulating sEV cargo, therapies that inhibit neutrophil recruitment or modulate sEVs release might reduce the pro-tumorigenic effects of 27HC. In addition, developing compounds that specifically block 27HC’s interaction with its receptors in neutrophils would decrease its ability to modulate sEVs cargo to promote cancer cell metastasis^17,19^. Moreover, combining 27HC pathway modulators, neutrophil recruitment inhibitors, and sEVs release modulators with existing breast cancer treatments, such as chemotherapy, might enhance overall efficacy. This approach could address the multiple mechanisms by which 27HC promotes tumor growth and metastasis. On the other hand, our data reveal an anti-cancer effect of neutrophils not exposed to 27HC; that their sEVs contain miRs that downregulate important pathways in cancer cells, such as WNT signaling. This speaks to the dual-nature of neutrophils (ie. pro- and anti-cancer), with their phenotype being context dependent. Regardless, neutrophils can still be harnessed as a way to reduce tumor progression.

Collectively, our results indicate that 27HC modulates miR levels in neutrophils, leading to the release of sEVs with modified miRs that are taken up by cancer cells. Ultimately leading to the upregulation of genes connected to WNT and stemness. Consequently, cancer cells lose their adherence in an EMT process, gaining migratory, stem-like phenotype, and chemoresistance.

## MATERIALS & METHODS

### REAGENTS

27-hydroxycholesterol (27HC, purity 95%) was synthesized and purchased from Sai Labs. Doxorubicin hydrochloride was purchased from MedChemExpress. Antibodies were purchased from BD Pharmingen: CD24 (FITC, Cat:561777), and CD44 (APC, Cat:559250). ALDH1A1 antibody stain was a gift from Professor Jefferson Chan Department of Chemistry, University of Illinois at Urbana-Champaign.

### ANIMALS

All protocols involving the use of animals were approved by the University of Illinois Institutional Animal Care and Use Committee (IACUC). Female mice between 8 and 10 weeks of age were purchased from Charles River Laboratories and housed at the University of Illinois in individually vented cages at 3 to 5 mice per cage. Food and water were provided ad libitum.

### CELL CULTURE

Medium for cell lines and primary cell culture - RPMI 1640 and DMEM, were supplemented with 1% penicillin/streptomycin (Corning), 1% nonessential amino acid (Corning), 1% sodium pyruvate (Corning), and 10% fetal bovine serum (FBS, Gibco). After adding mentioned reagents media was called “complete”. In experiments requiring the harvest or usage of EVs, 10% EV-depleted FBS (Gibco) was added in place of normal serum.

### CELL LINES

Murine mammary cancer cell lines - 4T1 and EMT6 were grown in Dulbecco’s modified Eagle’s medium supplemented with 1% nonessential amino acid, 1% sodium pyruvate, 1% penicillin/streptomycin, and 10% fetal bovine serum (FBS). In experiments requiring the harvest or usage of EVs, 10% EV-depleted FBS was added in place of normal serum.

### ISOLATION OF LY6G CELLS - NEUTROPHILS

Ly6G positive cells were isolated from mice bone marrow using Anti-Ly6G MicroBeads UltraPure (Miltenyi Biotec) according to the protocol. Briefly, femurs and tibia from 8–12-week-old WT BALB/c mice were collected and briefly soaked in 70% ethanol, and rinsed in phosphate-buffered saline (PBS, Lonza). Sterile medium (RPMI) was used to flush bone marrow over a 70-μM filter. Collected cells were used for Ly6G-positive cell isolation according to the protocol. Isolated cells were used immediately for experiments.

### EXTRACELLULAR VESICLES ISOLATION AND CHARACTERISATION

Extracellular vesicles were isolated from conditioned media after 24 and 48hrs of incubation using a commercially available kit - ExoQuict-TC ULTRA (SBI), according to the manufacturer’s protocol. Characterization of isolated SEVs was done previously, as of size characterization by Nanoparticle analysis (NanoSight NS300, Malvern Panalytical) and flow cytometry of known sEVs markers (CD9, CD63, CD81) and Ly6G using BD LSR Fortessa analyzer (BD Diagnostics)^44^. sEVs were stored at −80°C for no longer than 5 days after isolation prior to use.

### TREATMENT OF NEUTROPHILS WITH Let-7 INHIBITORS

After isolation, 0.45×10^6^ Ly6G positive cells were seeded in 6-well plates in 1.9 mL RPMI EVs depleted complete media and treated with four let-7 inhibitors or negative control for inhibitor (NC) cocktail (Qiagen miRCURY LNA miRNA inhibitors, a modified antisense). The transfection reaction was constructed according to the manufacturer’s protocol. Briefly, for 1 sample we prepared accordingly: 100uL of media, 3uL of HiPerFect transfection reagent (Qiagen), and 1uL of 10uM let-7 inhibitors (inhibitors for let-7a-5p, let-7g-5p, let7f-5p, and let7j) or NC. The final concentration of the inhibitors was 5nM. The prepared transfection reagent was mixed and incubated at room temperature for 10 min and added drop-wise to the cells. After 48h of culture cells were harvested. Cells were used for RNA and subsequent miR expression analysis as described below (see: *miRs QUANTIFICATION - REAL TIME-QUANTITATIVE POLYMERASE CHAIN REACTION (RT-qPCR)*). Conditioned media were collected for sEVs isolation followed by miRs expression evaluation and cancer cells treatments.

### CANCER CELL TREATMENT WITH SMALL EXTRACELLULAR VESICLES

20K of 4T1 and EMT6 cells were seeded in 12-well plates in 1mL of “complete” DMEM media supplemented with commercially available EV-depleted FBS (Gibco). At the time of seeding, sEVs were added to the cell culture, followed every 24h treatment.

### SMALL RNA SEQUENCING

#### Total RNA isolation

Total RNA from Ly6G cells and extracellular vesicles was isolated using miRNeasy Micro kit (Qiagen) according to the manufacturer’s protocol with supported modification for small RNA quantities. The quantity and quality of isolated RNA were determined with Qubit Fluorometer (Thermo Fisher Scientific), Bioanalyzer 2100 and Fragment Analyzer (Agilent).

#### Construction of miRNA libraries

Construction of libraries and sequencing on the Illumina NovaSeq 6000 were performed at the Roy J. Carver Biotechnology Center at the University of Illinois at Urbana-Champaign. Purified total RNAs from DMSO (vehicle) or 27HC treated cells’ EVs were converted into miRNA libraries with the QIAseq miRNA library preparation kit (Qiagen). The individually barcoded libraries were quantitated with Qubit (ThermoFisher) and run on a Fragment Analyzer (Agilent) to confirm the presence of a fragment of the expected length. The libraries were pooled in equimolar concentration and further quantitated by qPCR on a CFX Connect Real-Time qPCR system (Bio-Rad) for maximization of the number of clusters in the flow cell.

#### Sequencing of libraries in the NovaSeq

The barcoded miRNA libraries were loaded on a NovaSeq (Illumina) for cluster formation and sequencing. The libraries were sequenced from one end of the fragments for a total of 100 nucleotides. The fastq read files were generated and demultiplexed with the bcl2fastq v2.20 Conversion Software (Illumina). The quality of the demultiplexed fastq files was evaluated with the FastQC software, which generates reports with the quality scores, base composition, k-mer, GC and N contents, sequence duplication levels and overrepresented sequences.

#### Alignment and Counts

FASTQ read data were processed using the nf-core smrnaseq workflow v1.1.0^85^, using miRBase 22.1^86^ and Gencode M25^87^. The workflow was run using Nextflow v. 21.10.6^88^ with the command line call ‘nextflow run nf-core/smrnaseq -r 1.1.0 -c smrna.conf’, with the workflow configuration file settings including the miRNA protocol to ‘qiaseq’ and the species to ‘mmu’.

#### Statistical Analysis of small RNA-Seq

The mature counts were read into R (v4.1.2) and normalized using TMM^89^. miRNA genes without at least one count in four samples were removed. Final normalized expression values were made by re-running TMM and then converting to log2-based counts per million (logCPM) using edgeR’s cpm function^90^. Differential gene expression (DE) analysis was performed using the limma-trend method using a model of ∼Treatment + Group to control for batch effects of the sample processing group. Multiple testing correction was done using the False Discovery Rate method^91^. RNA-Seq data has been submitted to the NCBI Gene Expression Omnibus (accession number: <u>GSE272477).</u>

### DOWNSTREAM ANALYSIS OF SMALL RNS-SEQ

Different strategies were employed for downstream analysis. As the first step, we searched for targeted genes by our statistically differently expressed microRNA between DMSO vs. 27HC groups. Briefly, putative targeted genes were predicted by four different databases: TargetScanMouse (<u>TargetScanMouse 7.2</u>), miRDB (<u>miRDB - MicroRNA Target Prediction</u> <u>Database</u>), miRWalk (<u>Home - miRWalk (uni-heidelberg.de)</u>), and miRTarBase (<u>miRTarBase: the</u> <u>experimentally validated microRNA-target interactions database (cuhk.edu.cn)</u>). Only genes that were predicted to be potential targets of our microRNA in three out of four databases were taken under further investigation. In the first strategy, analysis of genes that were targeted by the greatest number of miRs were analyzed. In the second strategy, targeted genes were used for pathway analysis using four different approaches: KEGG (<u>KEGG: Kyoto Encyclopedia of Genes</u> <u>and Genomes</u>), PANTHERDB (pantherdb.org/?msclkid=3e954a57aba411ecb1b9fdc5b826ea61), g:Profiler (<u>g:Profiler – a web</u> <u>server for functional enrichment analysis and conversions of gene lists (ut.ee)</u>), and STRING (<u>STRING: functional protein association networks</u> (string-db.org)). The third strategy of downstream analysis employed mirPath v.3 software (<u>DIANA TOOLS - mirPath v.3 (uth.gr)</u>) for combined analysis of up and down-regulated microRNAs between tested groups, using microT-CDS (v5.0) as a database for pathway analysis.

### miR QUANTIFICATION - REAL TIME-QUANTITATIVE POLYMERASE CHAIN REACTION (RT-qPCR)

Isolated total RNA from Ly6G cells and extracellular vesicles was used for validation of microRNA sequencing by RT-qPCR according to the manufacturers’ protocols. Briefly, 10ng of isolated total RNA was reverse transcribed into complementary DNA (cDNA) using miRCURY LNA RT Kit (Qiagen). The cDNA was then amplified with specific commercially available primers for selected miRs using miRCURY LNA SYBR Green PCR Kit (Qiagen) using Bio-Rad CFX384 Thermal Cycler (Bio-Rad). In addition to each cDNA synthesis reaction, a UniSp6 RNA was added as an external control and used for data normalization and relative expression analysis via the 2^−ΔΔCT^ method.

### REAL TIME-QUANTITATIVE POLYMERASE CHAIN REACTION (RT-qPCR)

GeneJet RNA Purification kit (Thermo Fisher) was used for RNA extraction, according to the manufacturer’s protocols. cDNA was synthesized using iScript Reverse Transcription Supermix (Bio-Rad), according to the manufacturer protocols. Primer-BLAST software (https://www.ncbi.nlm.nih.gov/tools/primer-blast/) was used for primer design. A list of all used primers is presented in Supplementary Table 3. mRNA gene expression was quantified using iTaq Universal SYBR Green Supermix (Bio-Rad) on a Bio-Rad CFX384 Thermal Cycler (Bio-Rad). Relative expression was determined via the 2^−ΔΔCT^ method and normalized to the housekeeping gene (TATA box binding protein).

### FLOW CYTOMETRY

#### Cell death

For cell death assays, 20 × 10^5^ 4T1/EMT6 cells were seeded in 12 well plates and treated for 72/48hrs with sEVs from DMSO/27HC-neutrophils. Cells were harvested and stained using the <u>FITC</u> Annexin V/PI Dead Cell Apoptosis Kit (Invitrogen) following manufacturer protocol. Samples were analyzed using Attune Nxt Flow Cytometer. Cell status (live, early apoptotic, late apoptotic, or dead) was determined using FCS Express 6 Software.

#### Cancer stem cell markers

For CD44 and CD24 staining, cells were stained with fluorochrome-conjugated antibodies for cell surface antigens in FACS buffer (DPBS supplemented with 2% FBS and 1% penicillin/streptomycin) at 1:100 for 30 minutes at 4°C in the dark. Next, cells were fixed with 4% Formalin and incubated at 4°C in the dark until analysis. For the ALDH1A1 stain, cells were stained with AIDeSense live-cell dye^92^ for 30 minutes at 4°C in the dark with 2uM (dissolved in DMSO) dye. Samples were washed serially with FACS buffer and resuspended in FACS buffer for analysis. Cytometry data were acquired on either Attune Nxt Flow Cytometer. Cell status was determined using FCS Express 6 Software.

### TUMORSPHERE ASSAY

Tumorsphere assay was performed according to previously reported protocol^93^. Briefly, 1K of cells were seeded in ultra-low binding 24-well plates in 1mL 0.5% methylcellulose media supplemented with B27 (Gibco), epidermal growth factor (Corning), hydrocortisone (Sigma), insulin (Gibco), and B-ME (Sigma). On the seeding day, treatment with different doses of sEVs form DMSO or 27HC conditioned neutrophils were added. Sphere formation was evaluated under light microscope after 7 and 14 days and calculated using ImageJ software^94^.

### MIGRATION ASSAY

For migration assays, we used cancer cells after the sEVs treatment (see: *CANCER CELL TREATMENT WITH SMALL EXTRACELLULAR VESICLES*). Cells were seeded in EV depleted DMEM media on the top of 6.5mm 8um PET membrane in 24well plates for 24h. In the bottom of the wells was complete DMEM with normal (non-EV depleted) FBS. Migrated cells on the membrane were fixed and stained with 0.2% crystal violet according to the manufacturer’s protocol. Images were acquired at 2X magnification using an EVOS XL Digital Inverted Brightfield and Phase Contrast Microscope (Invitrogen, USA) and analyzed using ImageJ software^94^.

### 3D PROLIFERATION ASSAY

3D proliferation assay was performed as previously described^95^ and schematically presented in **Fig. 6G**. Briefly, 50µL of Matrigel was used to coat the wells in 96-well plate and incubated for 30 minutes in 37°C. 2000 cells were resuspended 100uL of complete DMEM with FBS EVs depleted <u>media</u>, supplemented with 2% Matrigel. After 24h, cells were treated daily with DMSO-sEVs and 27HC-sEVs. After three days, Doxorubicin was added at 2µM and proliferation was measured by measuring bioluminescence using Synergy LX Plate Reader (Agilent, USA).

### RNA-SEQ

Construction of the RNAseq libraries and sequencing on the Illumina NovaSeq 6000 were performed at the Roy J. Carver Biotechnology Center at the University of Illinois at Urbana-Champaign.

#### Construction of strand-specific RNAseq libraries

Total RNA was quantitated with Qubit high-sensitivity RNA reagent (Thermo Fisher) and the integrity and absence of genomic DNA was evaluated in a Fragment Analyzer (Agilent). The total RNAs were converted into individually barcoded polyadenylated mRNAseq libraries with the Kapa HyperPrep mRNA kit (Roche). Libraries were barcoded with Unique Dual Indexes (UDI’s) which have been developed to prevent index switching. The adaptor-ligated double-stranded cDNAs were amplified by PCR for 10 cycles with the Kapa HiFi polymerase (Roche). The final libraries were quantitated with Qubit (Thermo Fisher) and the average cDNA fragment sizes were determined on a Fragment Analyzer. The libraries were diluted to 5nM and further quantitated by qPCR on a CFX Connect Real-Time qPCR system (Bio-Rad) for accurate pooling of barcoded libraries and maximization of number of clusters in the flowcell.

#### Sequencing of libraries in the NovaSeq

The barcoded RNAseq libraries were loaded on one S1 lane on a NovaSeq 6000 for cluster formation and sequencing. The libraries were sequenced from both ends of the fragments for a total of 150bp from each end. The fastq read files were generated and demultiplexed with the bcl2fastq v2.20 Conversion Software (Illumina).

#### Quality check, Alignment and Counts of RNA-Seq

All reference files were downloaded from NCBI’s ftp site (https://ftp.ncbi.nlm.nih.gov/genomes/all/GCF/000/001/635/GCF_000001635.27_GRCm39/). The *Mus musculus* transcriptome file “GCF_000001635.27_GRCm39_rna.fna.gz” from Annotation 109 (https://www.ncbi.nlm.nih.gov/genome/annotation_euk/Mus_musculus/109/) from NCBI (https://www.ncbi.nlm.nih.gov/datasets/taxonomy/10090/) was used for quasi-mapping and count generation. This transcriptome is derived from genome GRCm39; Since the quasi-mapping step only uses transcript sequences, the gene model file “GGCF_000001635.27_GRCm39_genomic.gff.gz” was solely used to generate transcript-gene mapping table for obtaining gene-level counts.

The QC report was performed FASTQC (version 0.11.8, https://www.bioinformatics.babraham.ac.uk/projects/fastqc/) on individual samples then summarized into a single html report by using MultiQC^96^ (https://multiqc.info) version 1.11. The average per-base read quality scores are over 30 in all samples and no adapter sequences were found indicating those reads are high in quality. Thus, the trimming step was skipped, and sequences were directly proceeded to transcripts mapping and quantification.

Salmon version 1.10.0 was used to quasi-map reads to the transcriptome and quantify the abundance of each transcript^97^. The transcriptome was first indexed using the decoy-aware method in Salmon with the entire genome file “GCF_000001635.27_GRCm39_genomic.fna.gz” as the decoy sequence. Then quasi-mapping was performed to map reads to the transcriptome with additional arguments –seqBias and –gcBias to correct sequence-specific and GC content biases, --numBootstraps=30 to compute bootstrap transcript abundance estimates and – validateMappings and –recoverOrphans to help improve the accuracy of mappings. Gene-level counts were then estimated based on transcript-level counts using the “lengthScaledTPM” method from the tximport package. This method provides more accurate gene-level counts estimates and keeps multi-mapped reads in the analysis compared to traditional alignment-based method^98^. RNA-Seq data has been submitted to the NCBI Gene Expression Omnibus (accession number: GSE272476).

#### Statistical Analysis of RNA-Seq

When comparing expression levels, the numbers of reads per gene need to be normalized not only because of the differences in total number of reads, but because there could be differences in RNA composition such that the total number of reads would not be expected to be the same. The TMM (trimmed mean of M values) normalization^89^ in the edgeR package^90^ uses the assumption of *most genes do not change* to calculate a normalization factor for each sample to adjust for such biases in RNA composition.

Multidimensional scaling in the limma package^99^ was used as a sample QC step to check for outliers or batch effects. Differential gene expression (DE) analysis was performed using the limma-trend method^100^. First, an F-test was performed to identify genes that changed anywhere across any of the 6 treatments by comparing each of the groups back to D_A.48 as the baseline.

## Supporting information

Supplementary datasheet 1

Supplementary datasheet 2

Supplementary datasheet 3

## DOWNSTREAM ANALYSIS OF RNA-SEQ

Weighted gene co-expression network analysis (WGCNA) is a data mining method developed to discover the co-expressed gene clusters (modules) and detect the core genes (hub genes) in each module^101^. Specifically designed R package was used for this analysis^102^. The soft thresholding power 𝛽 was set to 5 to ensure a correlation coefficient close to 0.8. 16 modules were detected, and module heatmaps were plotted to visualize the expression pattern. Module eigengenes (ME) were calculated and visualized to represent the gene expression signatures of each module. Module membership (MM/kME) and gene significance (GS) were assessed for each module to understand the correlation of a gene to a module, and to assess the biological important of each gene accordingly. Upon identifying the module that shows significant different expression pattern between treatment groups, further analysis on gene enrichment, pathway, and clinical phenotype was performed using Shiny GO 0.80 version^67^.

## FIGURES DESCRIPTION

**Supplementary Figure 1.** *Gating strategy of cancer cells after Annexin V and PI staining for live-dead cells evaluation.* Gaiting strategy of **(A)** 4T1 after 72h and **(B)** EMT6 after 48h of exposure to DMSO-sEVs or 27HC-sEVs. Cells were stained with Annexin V (FITC – detection channel) and PI (PE-Texas Red – detection channel). Representative plot of no-stain, single stains with Annexin V and PI, and both stains of single cells suspension are presented.

**Supplementary Figure 2. *RNA characterization in sEVs from neutrophils treated with either DMSO or 27HC***. sEVs were isolated from the conditioned media of murine neutrophils treated with either DMSO or 27HC, followed by isolation of total RNA. (**A-B**) Representative Bioanalyzer 2000 results of total RNA isolated from DMSO-sEVs or 27HC-sEVs. (**C-D**) Representative Fragment Analyzer Agilent results of total RNA isolated from DMSO-sEVs or 27HC-sEVs. (**E)** Heatmap of all expressed miRs detected by small RNA-Seq, comparing expression between DMSO-sEVs and 27HC-sEVs. Each column represents an individual miR.

**Supplementary Figure 3. *Bioinformatic pipeline approach employed for miR target evaluation*.** Based on differently expressed miR cargo between 27HC-sEVs and DMSO-sEVs we employed different pipelines for target prediction pathways. 34 differentially expressed miRs were used in four different publicly available prediction models to predict their putative targets (TargetScanMouse, miRWalk, miRTarBas, and miRDB). The same software programs were then used for enrichment pathway analysis. As an additional approach for enrichment pathway analysis, we used DIANA mirPath v.3 software.

**Supplementary Figure 4.** *Expression of miRs in parental neutrophils and daughter sEVs after 24h and 48h of treatment with DMSO or 27HC.* Quantification of selected miRs in the parental neutrophils **(A, C)** and resulting sEVs **(B, D)**. **(A)** miR expression levels in neutrophils after 24h post treatment had shifted but were not yet statistically significant. **(B)** miR expression levels in sEVs 24h post treatment had not shifted. **(C)** miR expression levels in neutrophils after 48h post treatment had shifted in the expected direction with statistical significance. **(D)** miR expression levels in sEVs 48h post treatment had shifted in the expected direction with statistical significance. Statistical analyses were performed using one-way ANOVA followed by Tukey’s multiple comparison test, n=3; ****P-value<0.0001, ***P-value<0.001, **P-value<0.01. All data are presented as mean+/-SEM.

**Supplementary Figure 5. *Gene modules annotated by WGCNA analysis.*** Weighted gene co-expression network analysis (WGCNA) was utilized for bulk RNA-Seq analysis of 4T1 cells treated with DMSO-sEVs or 27HC-sEVs (data from Fig. 3). Heatmaps of the gene expression of the 16 detected modules is presented at the top, with the corresponding eigenvector graph below.

**Supplementary Figure 6.** *Expression of genes associated with the WNT pathway in cancer cells are regulated by 27HC-sEVs.* Expression of genes in the WNT pathway were measured by RT-qPCR through time post sEV-treatment. **(A)** 4T1 cells treated with sEVs. **(B)** EMT6 cells treated with sEVs. Statistical analyses were performed for each timepoint using one-way ANOVA followed by Tukey’s multiple comparison test n=6; ****P-value<0.0001, ***P-value<0.001, **P-value<0.01, *P-value<0.05. All data are presented as mean+/-SEM.

**Supplementary Figure 7.** Cancer cells treated with 27HC-sEVs adopt a *stem-like phenotype.* Expression of genes associated with pluripotency were measured by RT-qPCR in **(A)** 4T1 cells or **(B)** EMT6 cells after sEV treatment. Statistical analyses were performed using one-way ANOVA followed by Tukey’s multiple comparison test. n=3; ***P-value<0.001, **P-value<0.01, *P-value<0.05. ki67 expression in **(C)** 4T1 or **(D)** EMT6 cells after exposure to DMSO-sEVs or 27HC-sEVs. Statistical analyses were performed using one-way ANOVA followed by Tukey’s multiple comparison test. n=3; ****P-value<0.0001, ***P-value<0.001, **P-value<0.01, *P-value<0.05. **(E-F)** Flow cytometry quantification of ALDH1A1 positive 4T1 or EMT6 cells after exposure to DMSO-sEVs or 27HC-sEVs. Statistical analyses were performed using one-way ANOVA followed by Tukey’s multiple comparison test n=3; ****P-value<0.0001, ***P-value<0.001, **P-value<0.01, *P-value<0.05. All data are presented as mean+/-SEM. (**G**) Representative pictures of 4t1 tumorspheres after 7 days (first row) and 14 days (second raw) of culturing with different concentrations of DMSO-sEVs and 27HC-sEVs.

**Supplementary Table 1: *Pathways predicted to be downregulated by miRs in 27HC-sEVs.*** Three different pipelines were engaged for potential pathway analysis of detected differently expressed miRs in DMSO vs. 27HC – sEVs. A detailed description of the used methods is available in the Method sections and Supplementary Figure 3.

**Supplementary Table 2: *Overview of WGCNA statistical analysis between treatment groups in the Modules.*** Statistical differences in the Modules between the treatment groups in WGCNA of RNA-Seq. Modules are segregated based on statistical significance from the top to bottom. Da72 – 4T1 cells treated for 72h with DMSO-sEVs, adherent cells; Da48 – 4T1 cells treated for 48h with DMSO-sEVs, adherent cells; HCa48 – 4T1 cells treated for 48h with 27HC-sEVs, adherent cells; HCa72 – 4T1 cells treated for 72h with 27HC-sEVs, adherent cells; HCna48 – 4T1 cells treated for 48h with 27HC-sEVs, non-adherent cells; HCna72 – 4T1 cells treated for 72h with 27HC-sEVs, non-adherent cells.

**Supplementary Table 3:** *Sequence of primers used for gene expression analysis.* The sequence of forward and reverse primers used for RT-qPCR.

**Supplementary Datasheet 1**: ***RNA-Seq raw gene count and logarithmic counts per million reads.*** Raw gene counts (GeneLevel_counts), and logarithmic counts per million reads of the trimmed mean of M-values (TMMnormalized_logCPM). ENTREZID – gene identifier based on National Center for Biotechnology Information; symbol – gene symbol; product – full name of protein product of a given gene; D_A_48_1-5 - 4T1 cells treated for 48h with DMSO-sEVs, adherent cells, samples 1-5; D_A_72_1-5 - 4T1 cells treated for 72h with DMSO-sEVs, adherent cells, samples 1-5; HC_A_48_1-5 - 4T1 cells treated for 48h with 27HC-sEVs, adherent cells, samples 1-5; HC_A_72_1-5 - 4T1 cells treated for 72h with 27HC-sEVs, adherent cells, samples 1-5; HC_NA_48_1-5 - 4T1 cells treated for 48h with 27HC-sEVs, non-adherent cells, samples 1- 5; HC_NA_72_1-5 - 4T1 cells treated for 72h with 27HC-sEVs, non-adherent cells, samples 1-4.

**Supplementary Datasheet 2: *Gene segregation between the Modules in WGCNA.*** RNA-Seq genes selection for the created Modules in WGCNA analysis. SYMBOL – gene symbol; ENTREZID – gene identifier based on National Center for Biotechnology Information; logFC - logarithm of fold change; logCPM - logarithm of Counts Per Million; F – test value; FDR – false discovery rate; WGCNAmod – module number to which a given gene was segregated.

**Supplementary Datasheet 3: *List of HUB genes in Module 1 and Module 2 of WGCNA analysis.*** ENTREZID – gene identifier based on National Center for Biotechnology Information; kME - Module Eigengene; log10p-base-10 logarithm of the p-value; module – module selection; Symbol – gene symbol in the module.

## Funding

Department of Defense Era of Hope Scholar Award BC200206/W81XWH-20-BCRP-EOHS (ERN)

National Institutes of Health grant R01 CA234025 (ERN)

Postdoctoral Fellows Program at the Beckman Institute for Advanced Science and Technology (NK)

TiME Fellowship Award NIH T32EB019944 (CPS)

## Acknowledgments

The Cancer Center at Illinois Tumor Engineering and Phenotyping Shared Resource (TEP) provided certain cell lines and routine mycoplasma testing (directed and assisted by Hui Xu and Huimin Zhang). We would like to thank Chris Wright, an Associate Director of DNA Services at Roy J. Carver Biotechnology Center University of Illinois at Urbana-Champaign for her contribution to the experiment design and best practices for the sequencing experiments.

