## Supplementary Figures and Tables for "Neutrophils exposed to a cholesterol metabolite secrete extracellular vesicles that promote epithelial-mesenchymal transition and stemness in breast cancer cells"

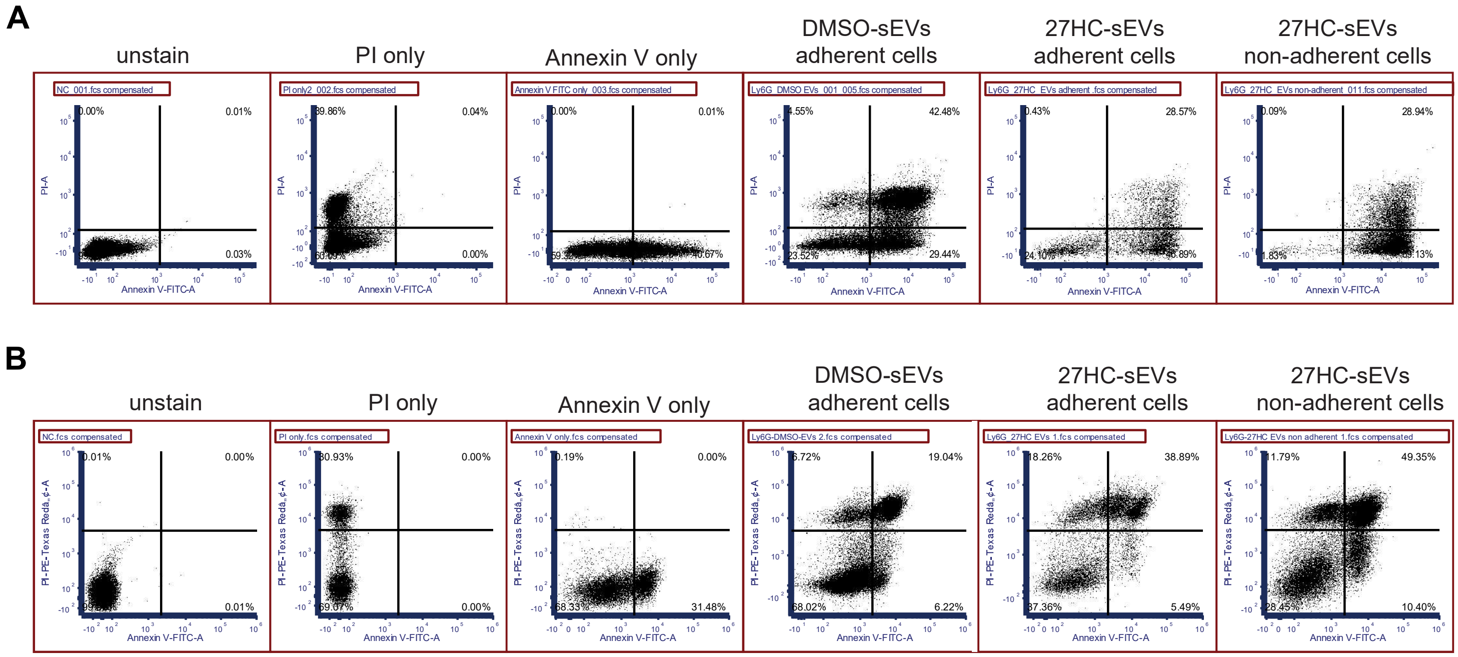


**Supplementary Figure 1.** ***Gating strategy of cancer cells after Annexin V and PI staining for live-dead cells evaluation.*** Gaiting strategy of **(A)** 4T1 after 72h and **(B)** EMT6 after 48h of exposure to DMSO-sEVs or 27HC-sEVs. Cells were stained with Annexin V (FITC – detection channel) and PI (PE-Texas Red – detection channel). Representative plot of no-stain, single stains with Annexin V and PI, and both stains of single cells suspension are presented.

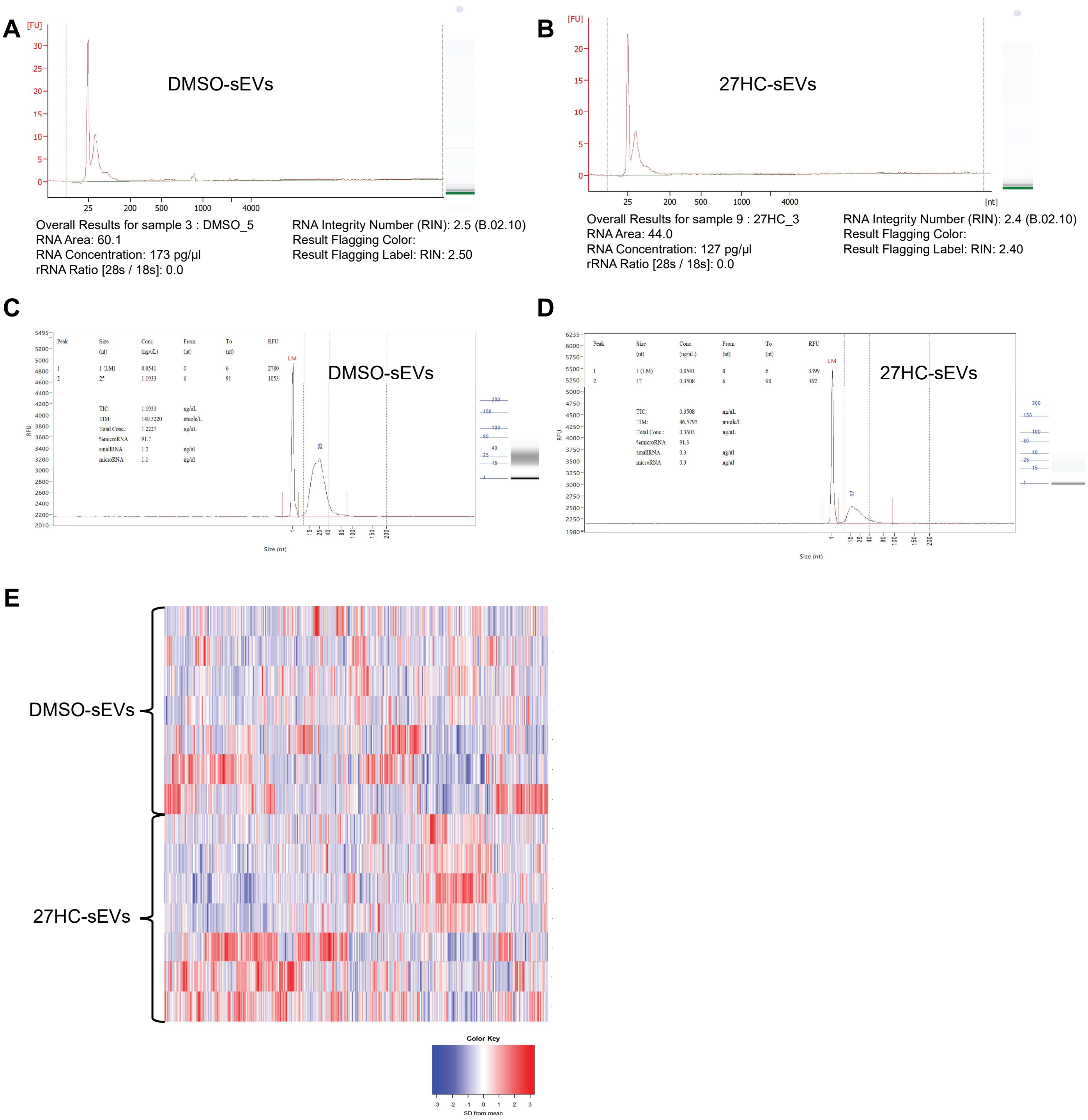


**Supplementary** ***Figure 2. RNA characterization in sEVs from neutrophils treated with either DMSO or 27HC***. sEVs were isolated from the conditioned media of murine neutrophils treated with either DMSO or 27HC, followed by isolation of total RNA. (**A-B**) Representative Bioanalyzer 2000 results of total RNA isolated from DMSO-sEVs or 27HC-sEVs. (**C-D**) Representative Fragment Analyzer Agilent results of total RNA isolated from DMSO-sEVs or 27HC-sEVs. (**E)** Heatmap of all expressed miRs detected by small RNA-Seq, comparing expression between DMSO-sEVs and 27HC-sEVs. Each column represents an individual miR.

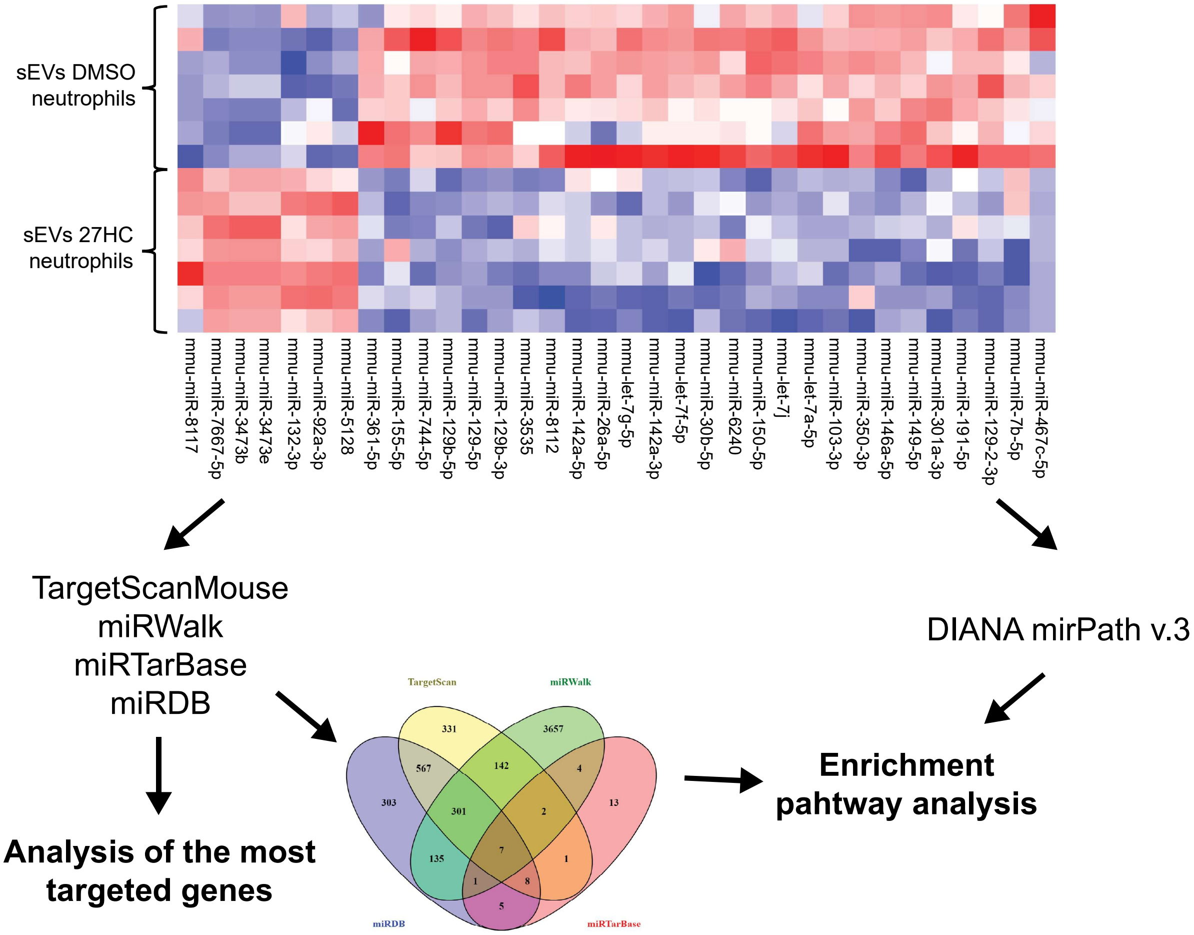


**Supplementary Figure 3.** ***Bioinformatic pipeline approach employed for miR target evaluation.*** Based on differently expressed miR cargo between 27HC-sEVs and DMSO-sEVs we employed different pipelines for target prediction pathways. 34 differentially expressed miRs were used in four different publicly available prediction models to predict their putative targets (TargetScanMouse, miRWalk, miRTarBas, and miRDB). The same software programs were then used for enrichment pathway analysis. As an additional approach for enrichment pathway analysis, we used DIANA mirPath v.3 software.

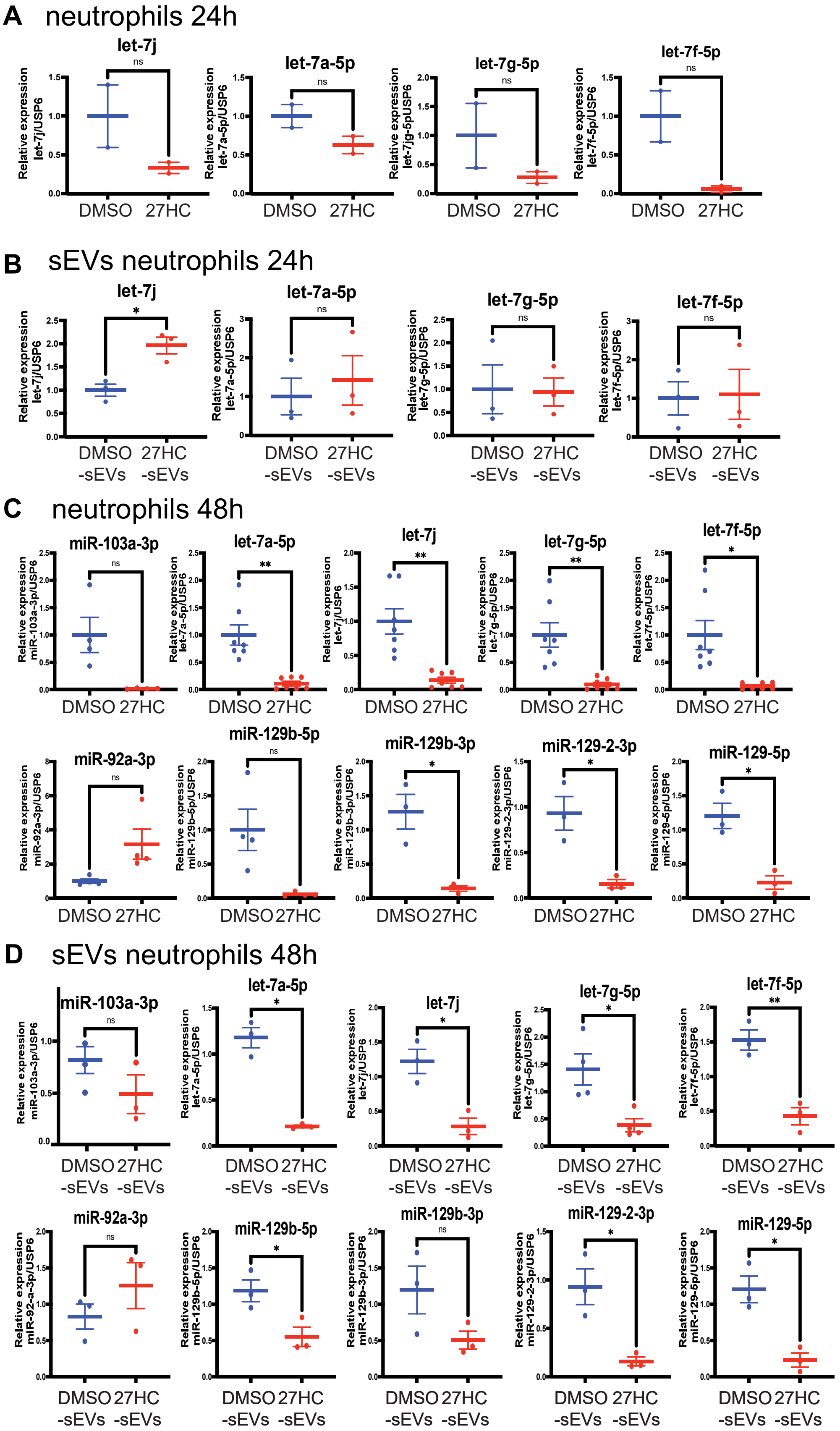


**Supplementary Figure 4.** ***Expression of miRs in parental neutrophils and daughter sEVs after 24h and 48h of treatment with DMSO or 27HC.*** Quantification of selected miRs in the parental neutrophils **(A, C)** and resulting sEVs **(B, D)**. **(A)** miR expression levels in neutrophils after 24h post treatment had shifted but were not yet statistically significant. **(B)** miR expression levels in sEVs 24h post treatment had not shifted. **(C)** miR expression levels in neutrophils after 48h post treatment had shifted in the expected direction with statistical significance. **(D)** miR expression levels in sEVs 48h post treatment had shifted in the expected direction with statistical significance. Statistical analyses were performed using one-way ANOVA followed by Tukey's multiple comparison test, n=3; ****P-value<0.0001, ***P-value<0.001, **P-value<0.01. All data are presented as mean+/-SEM.

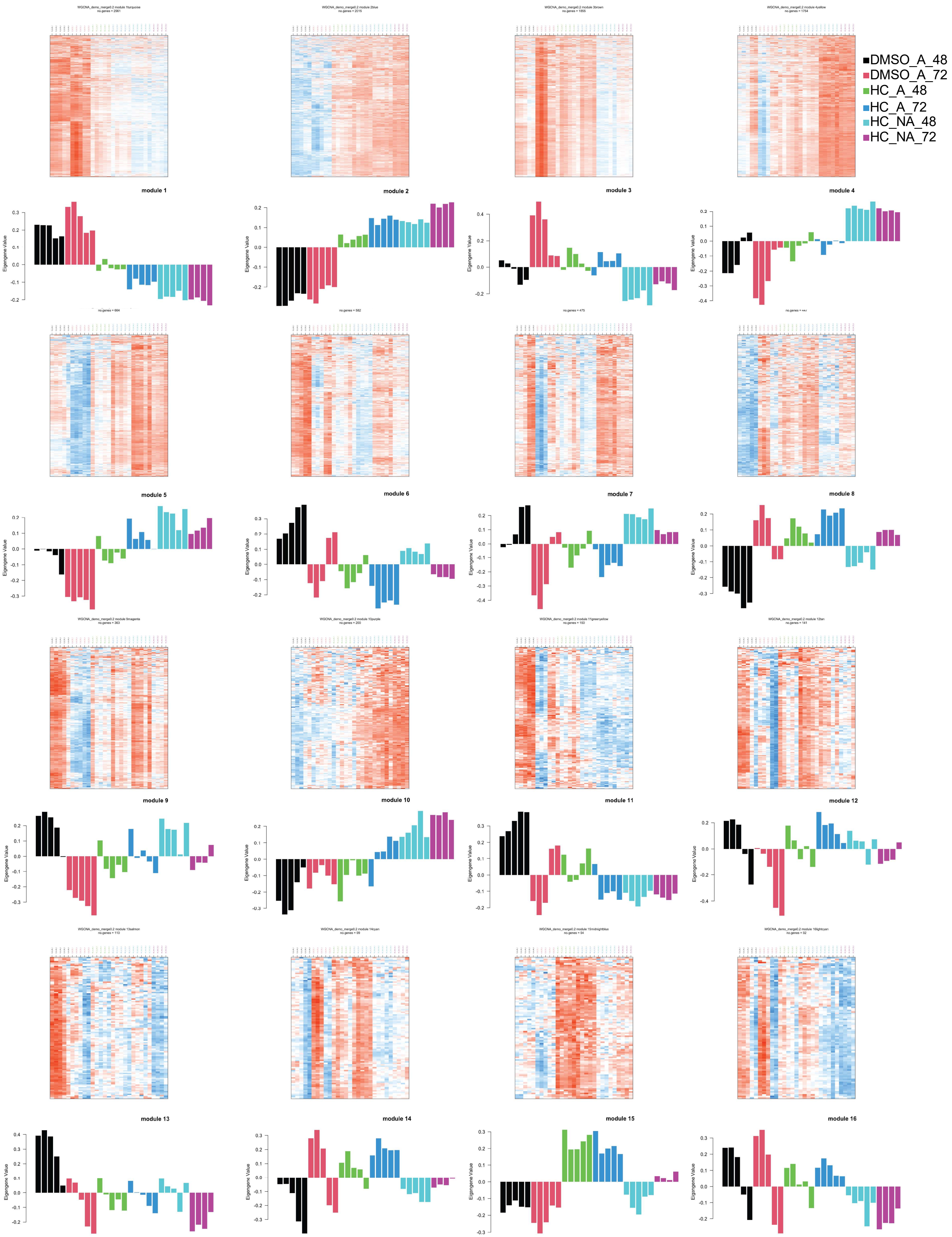


**Supplementary Figure 5**

**Supplementary Figure 5.** ***Gene modules annotated by WGCNA analysis.*** Weighted gene co-expression network analysis (WGCNA) was utilized for bulk RNA-Seq analysis of 4T1 cells treated with DMSO-sEVs or 27HC-sEVs (data from **Fig. 3**). Heatmaps of the gene expression of the 16 detected modules is presented at the top, with the corresponding eigenvector graph below.

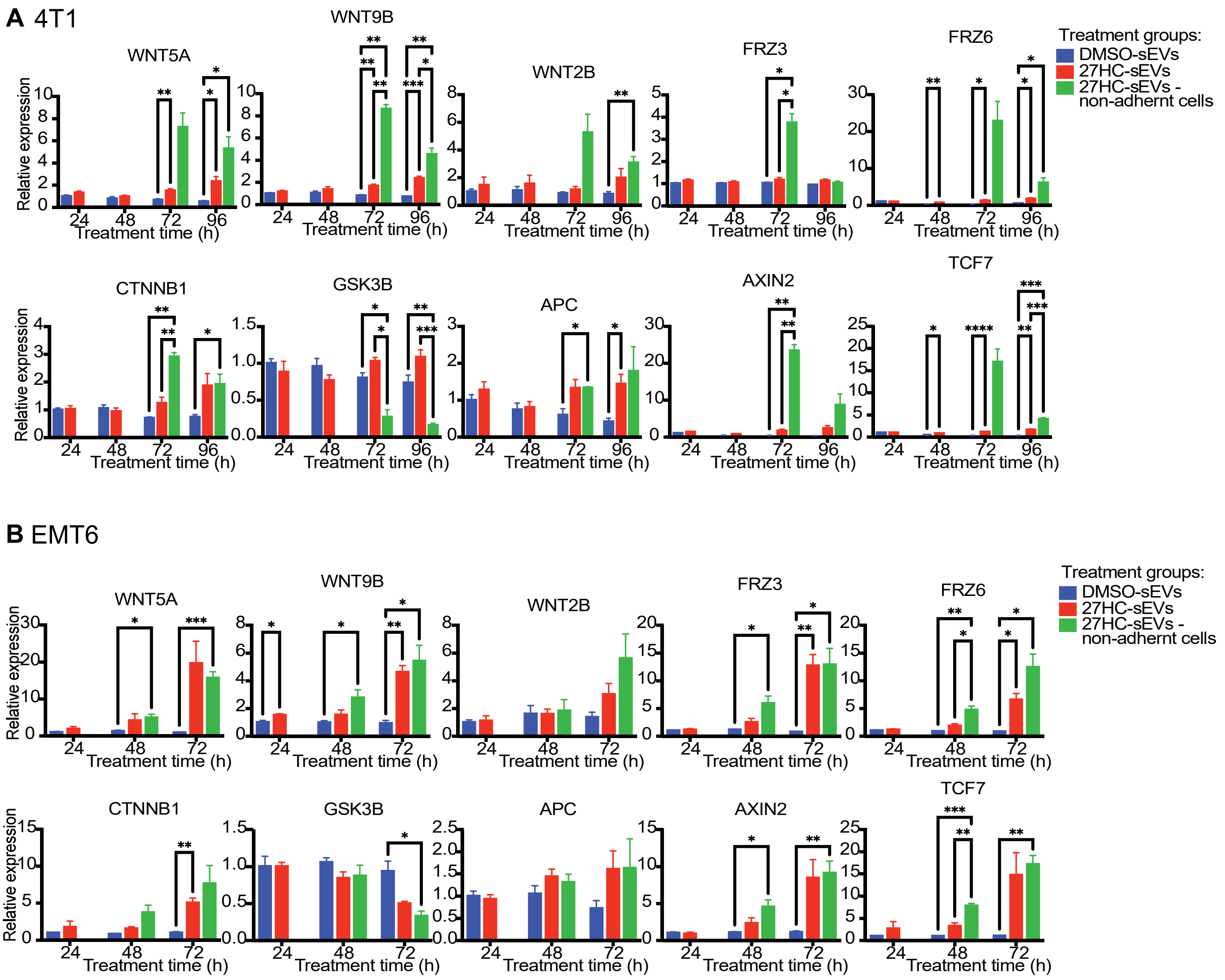


**Supplementary Figure 6. *Expression of genes associated with the WNT pathway in cancer cells are regulated by 27HC-sEVs.*** Expression of genes in the WNT pathway were measured by RT-qPCR through time post sEV-treatment. **(A)** 4T1 cells treated with sEVs. **(B)** EMT6 cells treated with sEVs. Statistical analyses were performed for each timepoint using one-way ANOVA followed by Tukey's multiple comparison test n=6; ****P-value<0.0001, ***P-value<0.001, **P-value<0.01, *P-value<0.05. All data are presented as mean+/-SEM.

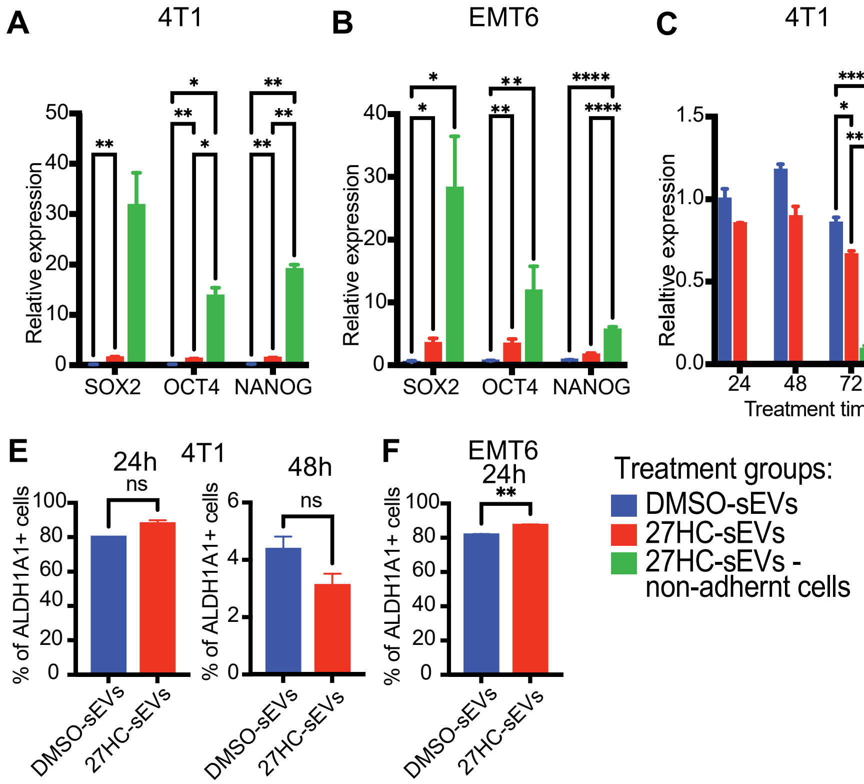


**Supplementary Figure 7. Cancer cells treated with 27HC-sEVs adopt a *stem-like phenotype.*** Expression of genes associated with pluripotency were measured by RT-qPCR in **(A)** 4T1 cells or **(B)** EMT6 cells after sEV treatment. Statistical analyses were performed using one-way ANOVA followed by Tukey's multiple comparison test. n=3; ***P-value<0.001, **P-value<0.01, *P-value<0.05. ki67 expression in **(C)** 4T1 or **(D)** EMT6 cells after exposure to DMSO-sEVs or 27HC-sEVs. Statistical analyses were performed using one-way ANOVA followed by Tukey's multiple comparison test. n=3; ****P-value<0.0001, ***P-value<0.001, **P-value<0.01, *P-value<0.05. **(E-F)** Flow cytometry quantification of ALDH1A1 positive 4T1 or EMT6 cells after exposure to DMSO-sEVs or 27HC-sEVs. Statistical analyses were performed using one-way ANOVA followed by Tukey's multiple comparison test n=3; ****P-value<0.0001, ***P-value<0.001, **P-value<0.01, *P-value<0.05. All data are presented as mean+/-SEM. (**G**) Representative pictures of 4t1 tumorspheres after 7 days (first row) and 14 days (second raw) of culturing with different concentrations of DMSO-sEVs and 27HC-sEVs.

**Supplementary Table 1: Pathways predicted to be by downregulated by miRs in 27HC-sEVs.** Three different pipelines were engaged for potential pathway analysis of detected differently expressed miRs in DMSO vs. 27HC – sEVs. A detailed description of used methods is available in the Method sections and Supplementary Figure 3.

| KEGG | PANTHERDB | g:profiler | STRING |
|---|---|---|---|
| ----- | Insulin/IGF pathway-protein kinase B signaling cascade | Insulin signaling (WP) | ----- |
| mmu04066 HIF-1 signaling pathway | Hypoxia response via HIF activation | GO:0071456 cellular response to hypoxia GO:0001666 response to hypoxia | GO:0071456 Cellular response to hypoxia<br>GO:0001666 Response to hypoxia<br>WP5024 Hypoxia-dependent proliferation of myoblasts |
| mmu04721 Synaptic vesicle cycle | Synaptic vesicle trafficking | GO:0008021 synaptic vesicle GO:0099504 synaptic vesicle cycle GO:0016079 synaptic vesicle exocytosis GO:2000300 regulation of synaptic vesicle exocytosis | GO:2000301 Negative regulation of synaptic vesicle exocytosis<br>GO:0099502 Calcium-dependent activation of synaptic vesicle fusion<br>GO:2000302 Positive regulation of synaptic vesicle exocytosis<br>GO:2000300 Regulation of synaptic vesicle exocytosis<br>GO:0098693 Regulation of synaptic vesicle cycle<br>GO:0016079 Synaptic vesicle exocytosis<br>GO:0099504 Synaptic vesicle cycle |
| mmu04151 PI3K-Akt signaling pathway | PI3 kinase pathway | KEGG:04151 PI3K-Akt signaling pathway | mmu04151 PI3K-Akt signaling pathway<br>MMU-199418 Negative regulation of the PI3K/AKT network<br>WP3855 Dysregulated miRNA targeting in insulin/PI3K-AKT signaling<br>WP2841 Focal adhesion: PI3K-Akt-mTOR signaling pathway |
| mmu04912 GnRH signaling pathway | Gonadotropin-releasing hormone receptor pathway | ----- | mmu04929 GnRH secretion<br>mmu04912 GnRH signaling pathway |
| mmu00190 Oxidative phosphorylation | Oxidative stress response | ----- | GO:1903203 Regulation of oxidative stress-induced neuron death<br>GO:1903204 Negative regulation of oxidative stress-induced neuron death |
| mmu04550 Signaling pathways regulating pluripotency of stem cells | Signaling pathways regulating pluripotency of stem cells | ----- | ----- |
| mmu04350 TGF-beta signaling pathway | TGF-beta signaling pathway | REAC:R-MMU-9006936 Signaling by TGFB family members | mmu04350 TGF-beta signaling pathway<br>WP113 TGF-beta signaling pathway |

| | | | |
|---|---|---|---|
| ----- | Angiogenesis | GO:0001525 angiogenesis<br>GO:0045765 regulation of angiogenesis<br>GO:0016525 negative regulation of angiogenesis<br>GO:1903670 regulation of sprouting angiogenesis | GO:1903672 Positive regulation of sprouting angiogenesis<br>GO:0090049 Regulation of cell migration involved in sprouting angiogenesis<br>GO:1903670 Regulation of sprouting angiogenesis<br>GO:0016525 Negative regulation of angiogenesis<br>GO:0045765 Regulation of angiogenesis<br>GO:0045766 Positive regulation of angiogenesis<br>GO:0001525 Angiogenesis |
| mmu05010 Alzheimer disease | Alzheimer disease-amyloid secretase pathway | GO:1904645 response to amyloid-beta<br>GO:1904646 cellular response to amyloid-beta | GO:1902004 Positive regulation of amyloid-beta formation<br>GO:1904646 Cellular response to amyloid-beta<br>GO:1902003 Regulation of amyloid-beta formation<br>GO:1902991 Regulation of amyloid precursor protein catabolic process |
| mmu01521 EGFR tyrosine kinase inhibitor resistance<br>mmu04370 VEGF signaling pathway | EGF receptor signaling pathway | KEGG:01521 EGFR tyrosine kinase inhibitor resistance | ----- |
| mmu04310 Wnt signaling pathway | Wnt signaling pathway | WP:WP539 Wnt signaling pathway (NetPath)<br>GO:0016055 Wnt signaling pathway<br>GO:0060070 canonical Wnt signaling pathway<br>GO:0030111 regulation of Wnt signaling pathway | WP539 Wnt signaling pathway (NetPath)<br>KW-0879 Wnt signaling pathway<br>GO:0060070 Canonical wnt signaling pathway<br>GO:0030111 Regulation of wnt signaling pathway<br>GO:0030178 Negative regulation of wnt signaling pathway<br>GO:0030177 Positive regulation of wnt signaling pathway<br>GO:0016055 Wnt signaling pathway<br>GO:0090090 Negative regulation of canonical wnt signaling pathway |
| mmu04010 MAPK signaling pathway | p38 MAPK pathway | KEGG:04010 MAPK signaling pathway | GO:0000165 MAPK cascade<br>mmu04010 MAPK signaling pathway<br>WP350 p38 Mapk signaling pathway |

**Supplementary Table 2:** ***Overview of WGCNA statistical analysis between treatment groups in the Modules.*** Statistical differences in the Modules between the treatment groups in WGCNA of RNA-Seq. Modules are segregated based on statistical significance from the top to bottom. Da72 – 4T1 cells treated for 72h with DMSO-sEVs, adherent cells; Da48 – 4T1 cells treated for 48h with DMSO-sEVs, adherent cells; HCa48 – 4T1 cells treated for 48h with 27HC-sEVs, adherent cells; HCa72 – 4T1 cells treated for 72h with 27HC-sEVs, adherent cells; HCna48 – 4T1 cells treated for 48h with 27HC-sEVs, non-adherent cells; HCna72 – 4T1 cells treated for 72h with 27HC-sEVs, non-adherent cells.

| Module | Da72_vs_Da48 | HCa48_vs_Da48 | HCa72_vs_Da48 | HCna48_vs_Da48 | HCna72_vs_Da48 | Interact | AveExpr | F | P.Value | adj.P.Val |
|---|---|---|---|---|---|---|---|---|---|---|
| ME2 | 0.03454782 | 0.31271926 | 0.40364059 | 0.39146276 | 0.47870636 | 0.27817144 | -4.22E-17 | 203.36573 | 4.00321314368437e-20 | 6.80546234426343e-19 |
| ME1 | 0.07027828 | -0.2154104 | -0.3094751 | -0.3823292 | -0.4055573 | -0.2856887 | -5.37E-17 | 104.271231 | 1.82999914347319e-16 | 1.55549927195221e-15 |
| ME15 | -0.0708837 | 0.39163063 | 0.35750624 | 0.02734448 | 0.17832797 | 0.4625143 | -5.50E-18 | 62.5124352 | 9.57380809384318e-14 | 5.42515791984447e-13 |
| ME5 | -0.2860356 | 0.01182618 | 0.12932771 | 0.26613649 | 0.1817376 | 0.29786183 | -1.84E-17 | 48.634538 | 1.88207466242431e-12 | 7.9988173153033e-12 |
| ME0 | 0.42859753 | 0.02610703 | 0.05833205 | 0.37824167 | 0.23119407 | -0.4024905 | -7.90E-17 | 33.3547986 | 1.4121071633113e-10 | 4.80116435525842e-10 |
| ME8 | 0.40292884 | 0.4064994 | 0.5050083 | 0.20739045 | 0.40761809 | 0.00357056 | 2.8951936633544e-17 | 24.5638802 | 3.93422611772092e-09 | 1.11469740002093e-08 |
| ME10 | 0.10870171 | 0.10902058 | 0.25436371 | 0.40535479 | 0.48341908 | 0.00031887 | -1.56E-17 | 21.8129015 | 1.36073048535445e-08 | 3.30463117871796e-08 |
| ME9 | -0.4970959 | -0.2547055 | -0.1856356 | -0.0326306 | -0.2227575 | 0.24239038 | -3.85E-17 | 19.6553579 | 3.93510969986932e-08 | 8.36210811222232e-08 |
| ME3 | 0.31588491 | 0.07698008 | 0.08105633 | -0.2060347 | -0.0997322 | -0.2389048 | -1.03E-17 | 17.3760053 | 1.33675105894906e-07 | 2.52497422245933e-07 |
| ME6 | -0.2945581 | -0.3447502 | -0.5175262 | -0.1840482 | -0.3630895 | -0.0501921 | 1.14850657719844e-17 | 15.8115111 | 3.32018565341731e-07 | 5.37888055709222e-07 |

| | | | | | | | | | | |
|---|---|---|---|---|---|---|---|---|---|---|
| ME4 | -0.134993 | 0.06711079 | 0.07826694 | 0.33217938 | 0.30762118 | 0.20210379 | 4.18726356270264e-17 | 15.7333276 | 3.48045212517732e-07 | 5.37888055709222e-07 |
| ME11 | -0.3679528 | -0.2643341 | -0.410705 | -0.459646 | -0.4525696 | 0.10361873 | -1.59E-17 | 14.5316137 | 7.34180829375959e-07 | 1.04008950828261e-06 |
| ME13 | -0.3792409 | -0.3411156 | -0.3331511 | -0.2809657 | -0.5156939 | 0.03812533 | -4.05E-17 | 10.8957633 | 9.41758084801018e-06 | 1.23152980320133e-05 |
| ME7 | -0.3106492 | -0.1585701 | -0.2589472 | 0.09233043 | -0.0311879 | 0.15207906 | 2.30897676457603e-17 | 7.76302246 | 0.00013558 | 0.00016463 |
| ME14 | 0.25839028 | 0.25210145 | 0.39074485 | 0.05190366 | 0.1371195 | -0.0062888 | 2.03381373045557e-18 | 5.64567332 | 0.00115921 | 0.00131377 |
| ME12 | -0.2887253 | -0.0542397 | 0.10053766 | -0.0206533 | -0.1246811 | 0.23448551 | -1.90E-17 | 3.88968697 | 0.00904214 | 0.00960728 |
| ME16 | -0.0145895 | -0.0478575 | 0.03036923 | -0.2003649 | -0.294456 | -0.033268 | 1.98595928973897e-17 | 3.02790576 | 0.02746514 | 0.02746514 |

**Supplementary table 3**. Sequence of primers used for gene expression analysis.

| Gene name | Sequence F (5'->3') | Sequence R (5'->3') |
|---|---|---|
| TBP | CAGGAGCCAAGAGTGAAGAACA | AAGAACTTAGCTGGGAAGCCC |
| SOX2 | GGAGGAGAGCGCCTGTTTTT | CTGGCGGAGAATAGTTGGGG |
| OCT4 | GGAGGGATGGCATACTGTGG | ACCTTTCCAAAGAGAACGCC |
| NANOG | TCGAATTCTGGGAACGCCTC | CAGGTCTTAACCTGCTTATAGCTCA |
| WNT2B | GACACGTCCTGGTGGTACATA | ACTGAGCGCATGATGTCTGG |
| WNT5A | GCTTTGGATTGTCCCCCAAG | ATTCCAATGGGCTTCTTCATGG |
| WNT9B | CTAGTGGCGCGAGGAGATG | GACCGGTCAGGCCGAAG |
| FRZ3 | GCAGATAGGTGGGCACAGTT | ATAGGGTGGAAGGGCTCCAT |
| FRZ6 | GCTCTCGCTCCCCGTTG | ATTCTGGGGCAACTGCTCG |
| APC | GGCTCGAAAATGGGGTCCA | TCCATCTTCAGTGCCTCAACT |
| GSK3B | GCTGTGTGTTGGCTGAATTGT | TGCTCCTGGTGAGTCCTTTGT |
| CTNNB1 | CGCCGCTTATAAATCGCTCC | TTCACAGGACACGAGCTGAC |
| AXIN2 | GCGCTTTGATAAGGTCCTGG | TCATGTGAGCCTCCTCTCTTTT |
| TCF7 | TAAACAGACCCCCGCCATCT | GGCTTGTTATGCAGCGGGG |
| ki67 | CCTGCCCGACCCTACAAAAT | TGCTGCTTCTCCTTCACTGG |
| SNAI1 | GGACGCGTGTGTGGAGTTCACC | TGGGCGGCTGGGAGCTTTTG |
| SLUG | AGCAGCTGCACTGTGATGCCC | GCAAGAGAAAGGCTTTTCCCCAGTG |
| CDH-1 | CTGGGCTGGACCGAGAGAGTT | TTGAGTGTGGCGATCCGGGC |
| CDH-2 | GCCTTGCTTCAGGCGTCTGTG | TGCCGTCCTCGTCCACCTTGA |
| VIM | AACAACGATGCCCTGCGCCA | GCTCCAGGGACTCGTTAGTGCCTTT |
| TWIST1 | GGACAAGCTGAGCAAGATTCA | CGGAGAAGGCGTAGCTGAG |
